# Temperature Dramatically Shapes Mosquito Gene Expression with Consequences for Mosquito-Zika Virus Interactions

**DOI:** 10.1101/783035

**Authors:** Priscila Gonçalves Ferreira, Blanka Tesla, Elvira Cynthia Alves Horácio, Laila Alves Nahum, Melinda Ann Brindley, Tiago Antônio de Oliveira Mendes, Courtney Cuinn Murdock

**Affiliations:** Department of Biochemistry and Molecular Biology, Universidade Federal de Viçosa, Viçosa, Minas Gerais, Brazil; Department of Infectious Diseases, College of Veterinary Medicine, University of Georgia, Athens, Georgia, United States of America; René Rachou Institute, Oswaldo Cruz Foundation, Belo Horizonte, Minas Gerais, Brazil; Department of Genetics, Ecology and Evolution, Biological Sciences Institute, Federal University of Minas Gerais, Belo Horizonte, Minas Gerais, Brazil; Promove College of Technology, Belo Horizonte, Minas Gerais, Brazil; Department of Population Health, College of Veterinary Medicine, University of Georgia, Athens, GA, United States of America; Center for Vaccines and Immunology, University of Georgia, Athens GA, United States of America; Odum School of Ecology, University of Georgia, Athens GA United States of America; Center for Ecology of Infectious Diseases, University of Georgia, Athens GA, United States of America; Center for Emerging and Global Tropical Diseases, University of Georgia, Athens GA, United States of America; Riverbasin Center, University of Georgia, Athens GA, United States of America

**Keywords:** Temperature, *Aedes aegypti*, Zika virus, RNA-seq, Transcriptome, Immune response

## Abstract

Vector-borne flaviviruses are emerging threats to human health. For successful transmission, the virus needs to efficiently enter mosquito cells, replicate within, and escape several tissue barriers while mosquitoes elicit major transcriptional responses to flavivirus infection. This process will not only be affected by the specific mosquito-pathogen pairing, but also variation in key environmental variables such as temperature. Thus far, few studies have examined the molecular responses triggered by temperature and how these responses modify infection outcomes despite substantial evidence showing strong relationships between temperature and transmission in a diversity of systems. To define the host transcriptional changes associated with temperature variation during the early infection process, we compared the transcriptome of mosquito midgut samples from mosquitoes exposed to Zika virus (ZIKV) and non-exposed mosquitoes housed at three different temperatures (20, 28, and 36°C). While the high temperature samples did not have significant changes from standard rearing conditions (28°C) 48 hr post-exposure, the transcriptome profile of mosquitoes housed at 20°C was dramatically different. The expression of genes most altered by the cooler temperature involved aspects of blood-meal digestion, ROS metabolism, and mosquito innate immunity. Further, we did not find significant differences in the viral RNA copy number between 24 and 48 hr post-exposure at 20°C, suggesting ZIKV replication is limited by cold-induced changes to the mosquito midgut environment. In ZIKV-exposed mosquitoes, vitellogenin, a lipid carrier protein, was the most up-regulated at 20°C. Our results provide a deeper understanding of the temperature-triggered transcriptional changes in *Aedes aegypti* and can be used to further define the molecular mechanisms driven by environmental temperature variation.

**Contribution to the Field Statement:** A variety of methods for engineering refractory mosquitoes are currently being studied and show promise for disease control. Although considerable effort has been put into understanding the immune system of mosquitoes in response to infections, almost nothing is understood about environmental influences in regulating these responses. Here, we used RNA sequencing to study the effect of temperature on the mosquito transcriptome profile, as well as assess the changes in the immune response to ZIKV infection at three different temperatures. We found a remarkable effect of temperature on the transcriptome profile of mosquitoes exposed to cool conditions (20°C) after imbibing a blood meal, as well as accumulation of transcripts involved with different mechanisms associated with blood meal digestion, metabolism, and some components of the immune response in mosquitoes. Our results provide new insights in potential mechanisms that limit temperature-driven pathogen establishment and replication within the mosquito vector.

## Introduction

Over the past three decades, there have been significant advances in our understanding of the physiological and molecular interactions between pathogens and mosquito vectors (Bartholomay and Michel, 2018; Kumar et al., 2018; Simões et al., 2018). Research has provided substantial insights into the immune genes and pathways that shape resistance to vector-borne pathogens and has revealed many promising targets for genetic manipulation (Wilke and Marrelli, 2015; Yen et al., 2018; Shaw and Catteruccia, 2019). Yet, we are only just beginning to understand the complexity underlying mosquito-pathogen interactions. The response mosquitoes mount toward a given pathogen is a dynamic phenotype, which is dependent upon both the specific mosquito pathogen pairing (Lambrechts et al., 2013; Zouache et al., 2014; Duggal et al., 2015; Chouin-Carneiro et al., 2016) as well as variation in key environmental factors (Okech et al., 2007; Alto et al., 2008; Parham and Michael, 2010; Parham et al., 2015; Cohen et al., 2016; Gloria-Soria et al., 2017; Murdock et al., 2017; Shragai et al., 2017; Siraj et al., 2018). Knowledge of how vectors respond to environmental variation is especially relevant for understanding how vector-borne pathogens emerge and defining the biological constraints on transmission.

Vector-borne flaviviruses are emerging threats to human health. Yellow fever virus (YFV), dengue virus (DENV), West Nile virus (WNV), and most recently, Zika virus (ZIKV) can be found throughout tropical and subtropical zones. For successful transmission, flaviviruses are taken up through the bite of a mosquito vector when it takes a blood meal from an infectious host. To complete infection within the mosquito vector, the virus needs to efficiently enter, replicate within, and escape several tissue barriers, primarily the midgut and salivary glands (Franz et al., 2015; Kumar et al., 2018). A number of studies have demonstrated that mosquitoes elicit major transcriptional changes in response to flavivirus infection, which could play an important role in limiting infection (Xi et al., 2008; Sim and Dimopoulos, 2010; Colpitts et al., 2011; Bonizzoni et al., 2012; Chauhan et al., 2012; Etebari et al., 2017; Saldaña et al., 2017a). These include differential regulation of genes involved in RNA interference (RNAi), classical immune pathways (e.g. JAK-STAT, Toll), production and transport of energy, metabolism, as well as the production of noncoding RNAs (small and long) and microRNAs that could be involved in targeted gene regulation. Currently, it is unclear how conserved these responses to infection are across different mosquito-flavivirus combinations (Etebari et al., 2017) and how relevant environmental variation shapes the nature, magnitude, and timing of these responses (Murdock et al., 2012b, 2014; Adelman et al., 2013).

One of the major environmental variables that influence mosquito-pathogen interactions, as well as vector-borne transmission, is variation in environmental temperature. Mosquitoes are small ectothermic organisms, and many studies have already demonstrated that temperature can markedly affect diverse aspects of mosquito physiology and ecology, as well as pathogen replication. These combined effects result in temperature significantly impacting the proportion of the mosquito population that becomes infected, goes on to become infectious, and overall transmission potential (Christofferson and Mores, 2016; Tesla et al., 2018a; Mordecai et al., 2019). The extent to which temperature shapes transmission directly, through effects on pathogen biology, or indirectly, through effects on mosquito immunity and physiology, remains largely unexplored (Murdock et al., 2012a).

In previous work (Tesla et al., 2018a), we have demonstrated that ZIKV transmission by *Aedes aegypti* is optimized at a mean temperature of 29°C and has a thermal range from 22.7°C to 34.7°C. Further, we observed constraints on ZIKV transmission at cool and warm temperatures to be mediated by different factors. Cool temperatures inhibited ZIKV transmission due to slow virus replication and escape from the midgut resulting in decreased vector competence and long extrinsic incubation periods. In contrast, increased mosquito mortality at hotter temperatures constrained transmission, despite efficient ZIKV infection and rapid dissemination.

These temperature constraints on infection are likely regulated through different mechanisms. Temperature variation in general could profoundly impact arbovirus infection and replication early in infection due to shifts in the balance and dynamics of the midgut environment, the first host environment encountered. This environment is fairly complex as arboviruses encounter microbiota (Xi et al., 2008; Carissimo et al., 2014; Hegde et al., 2015; Saraiva et al., 2016; Barletta et al., 2017; Saldaña et al., 2017b), oxidative and nitration stress associated with digestion of the blood meal (Luckhart et al., 1998; Graça-Souza et al., 2006; Xi et al., 2008), and key immune factors (Campbell et al., 2008; Xi et al., 2008; Cirimotich et al., 2009; Sánchez-Vargas et al., 2009; Souza-Neto et al., 2009). To better understand the effects of temperature on the ZIKV-mosquito interaction, we used RNA sequencing to describe the transcriptional response of *Ae. aegypti* midguts to ZIKV infection at three different temperatures (20°C, 28°C, and 36°C) early on in the infection process.

## Methods

### Ethics statement

All mosquito infection work with ZIKV was approved by the University of Georgia, Athens Institutional Biosafety Committee (reference number 2015-0039).

### Virus production

ZIKV MEX1-44 was isolated from *Ae. aegypti* mosquitoes from Tapachula, Chiapas, Mexico in January 2016, kindly provided by University of Texas Medical Branch Arbovirus Reference Collection. ZIKV stocks were propagated in Vero cells cultured in DMEM (Dulbeco’s Modified Eagle Medium), 5% fetal bovine serum (FBS) at 37°C and 5% CO_2_. Four days following inoculation, when cells showed visible cytopathic effect (> 90%), supernatant containing virus was collected, cell debris was cleared by centrifugation (3,500xg for 5 min), and virus was aliquoted and frozen at −80°C. The stock was titrated using standard plaque assays on Vero cells (Willard et al., 2017) and expressed in plaque forming units per millilitre (PFU/mL).

### Mosquito husbandry

*Ae. aegypti* eggs collected in Chiapas, Mexico, were hatched in ddH_2_O under reduced pressure in a vacuum desiccator. Larvae were grown in trays with 200 larvae in 1L ddH_2_O and 4 fish food pellets (Hikari Cichlid Cod Fish Pellets). Emerging adults were kept in rearing cages and fed with 10% sucrose solution *ad libitum*. Colonies were maintained on O-positive human blood (Interstate Blood Bank, male donors between 30-35 yr). Both, larvae and adults were maintained under standard insectary conditions (27°C ± 0.5°C, 80% ± 5% relative humidity, and a 12:12 light: dark diurnal cycle) (Percival Scientific).

### Experimental mosquito infection

Briefly, 3 to 4-day-old female mosquitoes (F5 generation; n = 600) were separated using a vacuum aspirator, transferred to an arthropod containment level three (ACL-3) facility at the University of Georgia and housed at 28°C ± 0.5°C, 80% ± 5% relative humidity, 12 hr:12 hr light:dark cycle (Percival Scientific). Mosquitoes were fed with a non-infectious or ZIKV-containing blood-meal (10^6^ PFU/mL) after a 12 hr period of starvation in a manner previously described (Tesla et al., 2018b). After the feed, engorged ZIKV-exposed (n = 120) and unexposed mosquitoes (n = 120) were randomly allocated across 12 paper cups (6 with ZIKV-exposed and 6 with ZIKV-unexposed mosquitoes, n = 20 mosquitoes per cup). Forty engorged females that fed on either infectious blood or non-infectious blood were randomly distributed across one of three temperature treatments, 20°C, 28°C, and 36°C (± 0.5°C); at 80% ± 5% relative humidity; and 12 hr:12 hr light:dark cycle. Mosquitoes were provided 10% sucrose *ad libitum* throughout the duration of the experiment (48 hr), and three full biological replicates were performed.

### Viral genome quantificantion

Viral genomes were detected as previously described (Willard et al., 2019). Briefly dissected mosquito midguts were collected in pools of 5 and viral RNA was isolated using the Zymo Quick-RNA™ Viral Kit (Zymo, Irvine, CA, USA). Viral RNA samples were reverse-transcribed (RT) to cDNA (High Capacity RNA-to-cDNA Kit, Applied Biosystems, Foster City, CA, USA). To quantify the number of ZIKV genomes we used the cDNA in a quantitative PCR (qPCR) reaction assay using TaqMan Gene Expression Master Mix (Applied Biosystems, ThermoFisher, Waltham, MA, USA), primers and probes (F: ZIKV 1086, R: ZIKV 1162c, ZIKV 1107-FAM; TaqMan MGB Probe; Invitrogen Custom Primers) (Lanciotti et al., 2008). Each sample was analyzed in duplicate, and each plate contained a DNA plasmid standard curve (ZIKV molecular clone), no template, and no primer controls. ZIKV copy numbers were extrapolated from the generated standard curve using the Applied Biosystems protocol. The limit of detection was experimentally established to be 30 copies. Final copy numbers were adjusted by back-calculations to the total RNA and cDNA volume and expressed as copies per 5 midgut pool.

### Midgut dissection and RNA isolation

Midgut immune responses contribute significantly to viral bottlenecks during infection and these processes could dynamically differ with temperature. To capture gene expression changes in the mosquito midgut early during the infection process, we isolated total RNA from midgut tissues. Fifteen ZIKV-exposed and 15 non-exposed mosquitoes were removed from each temperature at 24 and 48 hours post-infection for RNA sequencing. The mosquitoes were killed with cold ethanol and washed in cold PBS containing 0.2% RNase inhibitor (Sigma-Aldrich) and 0.1% DEPC (Amresco). Midguts were carefully dissected and stored immediately in RNA*later* (Invitrogen) at 4°C for 24 hr after which they were transferred to −80°C. Total RNA was isolated from pools of 15 midguts using the Qiagen RNeasy Mini Kit as per the manufacturer’s instructions.

### Library preparation and sequencing

Extracted RNA was sent to University of Georgia Genomics Core for cDNA library preparation and RNA sequencing. The quality of the total RNA was analysed using an Agilent Bioanalyzer RNA 6000 Pico kit (Agilent Technnology). Poly(A) mRNA from total RNA were captured by magnetic oligo-dT beads and fragmented by heating (94°C, 6 min) in the presence of divalent cations (Mg^2+^) using the KAPA Stranded mRNA-Seq Kit for Illumina®. Fragmented poly(A) mRNA samples were converted to cDNA using random priming. After second-strand synthesis, an adenine residue was added to 3’-end of dscDNA fragments and the Illumina TruSeqLT adapter ligation to 3’-dAMP library fragments was performed. Adapter-ligated DNA was amplified by PCR using the KAPA Library Amplification Primer Mix. After creating the library of DNA templates, the fragment size distribution of these libraries was confirmed by electrophoresis and the library concentration was determined by qPCR. Sequencing was completed on an Illumina NextSeq 500 using the PE75bp (75bp sequencing reads) settings on a High Output 150 cycle kit using Illumina standard protocols for loading. Raw sequencing files were demultiplexed using the Illumina BaseSpace cloud platform demultiplexing pipeline. Four technical replicates were run per sample.

### RNA sequence analysis in response to temperature and Zika virus exposure

The Tuxedo suite of tools was used to analyse the RNA-seq data (Trapnell et al., 2013). At each stage of the analysis pipeline, we were careful to identify and correct possible sources of bias in the study (Conesa et al., 2016). Read quality was assessed using FastQC (Andrews, 2010). Poor quality reads (quality score below 20), short reads (less than 25 bases), and adapter sequences (Bolger et al., 2014) were removed using Trimmomatic (v 0.36).

We aligned and mapped clean reads up to two different loci to the *Ae. aegypti* genome (NCBI ID: GCA_002204515.1) obtained from NCBI (https://www.ncbi.nlm.nih.gov/) using TopHat (v 2.1.1) (Trapnell et al., 2009). We then performed differential gene expression analysis on RNAseq reads using Cuffdiff (Trapnell et al., 2013), which calculates expression levels in response to experimental treatments of interest (e.g. temperature and ZIKV-exposure) and determines if observed variation in expression levels across groups are significantly different. The *Ae. aegypti* genome and its annotation file (NCBI ID: GCA_002204515.1) were used to run bias detection and determine the transcripts examined for differential expression, respectively. The relative expression levels were produced as fragments per kilobase of transcript per million fragments mapped (FPKM values), which normalized read count based on gene length and the total number of mapped reads. After quantification and normalization of expression values, differential expression analysis was carried out on the experimental data from 24 and 48 hr following the blood meal. To characterize changes in the RNA transcriptome in response to temperature, we compared the RNA expression profiles of unexposed mosquitoes maintained at standard conditions and near the predicted thermal optimum for ZIKV transmission (28°C) to those housed at cool (20°C) and hot (36°C) conditions. To determine if temperature modified global RNA expression in ZIKV-exposed individuals, we compared whether the variation in expression profiles of ZIKV-exposed individuals across temperature treatments was similar to their unexposed counterparts. Finally, to evaluate the effects of ZIKV infection on global RNA expression, we compared RNA expression between unexposed and ZIKV-exposed mosquitoes within a given temperature treatment. The data quality assessment from the RNAseq analysis was performed using R package (v 3.4.4) (Team, 2008) and plotted using GraphPad Prism (version 5.01 for Windows, GraphPad Software, San Diego California USA, www.graphpad.com). Perl scripts were written to extract specific information from RNAseq analysis output files necessary for each assessment of the results obtained. Principal-component analysis (PCA) was employed to examine the quality of replicates, the overall differences between samples and to determine the proximity among the experimental groups.

### Gene Ontology (GO) analysis

We employed BiNGO (v 3.0.3) (Maere et al., 2005) to determine which GO terms were significantly over-represented in a set of genes. First, the FASTA sequences of all differentially expressed genes were recovered using the protein database on the NCBI Batch Entrez (https://www.ncbi.nlm.nih.gov/sites/batchentrez). The sequences were input into STRING (v 10.5) (Szklarczyk et al., 2017) to predict protein-protein interactions. GO functional annotations were provided by AgBase-Goanna (v 2.00) (Mccarthy et al., 2006) using the *Ae. aegypti* protein sequences in FASTA format from the STRING database as an input file. The similarity search for GO annotations were performed by Blastp using UniProt Database, E-value: 10e^-5^, Matrix: BLOSUM62 and word size: 3. This file annotation was employed as a reference set for an enrichment analysis. We used the default mode employing Hypergeometric tests for assessing the enrichment of a GO term in the test set, and *p*-values were corrected by Benjamini & Hochberg False Discovery Rate (FDR) correction to control for type I error. Corrected *p*-values less than 0.05 were considered significant.

## Results

From a total of 36 RNA samples that were sequenced using the Illumina platform, the number of clean paired reads varied between 2,063,917 and 3,028,305 reads for each library. All libraries resulted in a concordant pair alignment rate higher than 70%.

### Cool temperatures restrict ZIKV replication in the mosquito midgut

Viral RNA quantification of exposed mosquitoes revealed levels of ZIKV RNA in midgut samples from mosquitoes that imbibed a blood meal containing ZIKV (Figure 1). ZIKV replication was evident in mosquitoes housed at 28°C and 36°C, with increases in mosquito RNA copy number between 24 and 48 hr post-feed (hpf). Further, the efficiency of ZIKV replication was maximized at the warmest temperature with mosquitoes housed at 36°C having higher ZIKV RNA levels in their midguts relative to those housed at 28°C. Although the presence of ZIKV RNA was observed 24 and 48 hr following the infectious blood meal in mosquitoes housed in the cool environment (20°C), we did not find amplification of viral genomes (Figure 1). This result suggests that cooler temperatures constrain ZIKV replication in the midgut.

**Figure 1.**
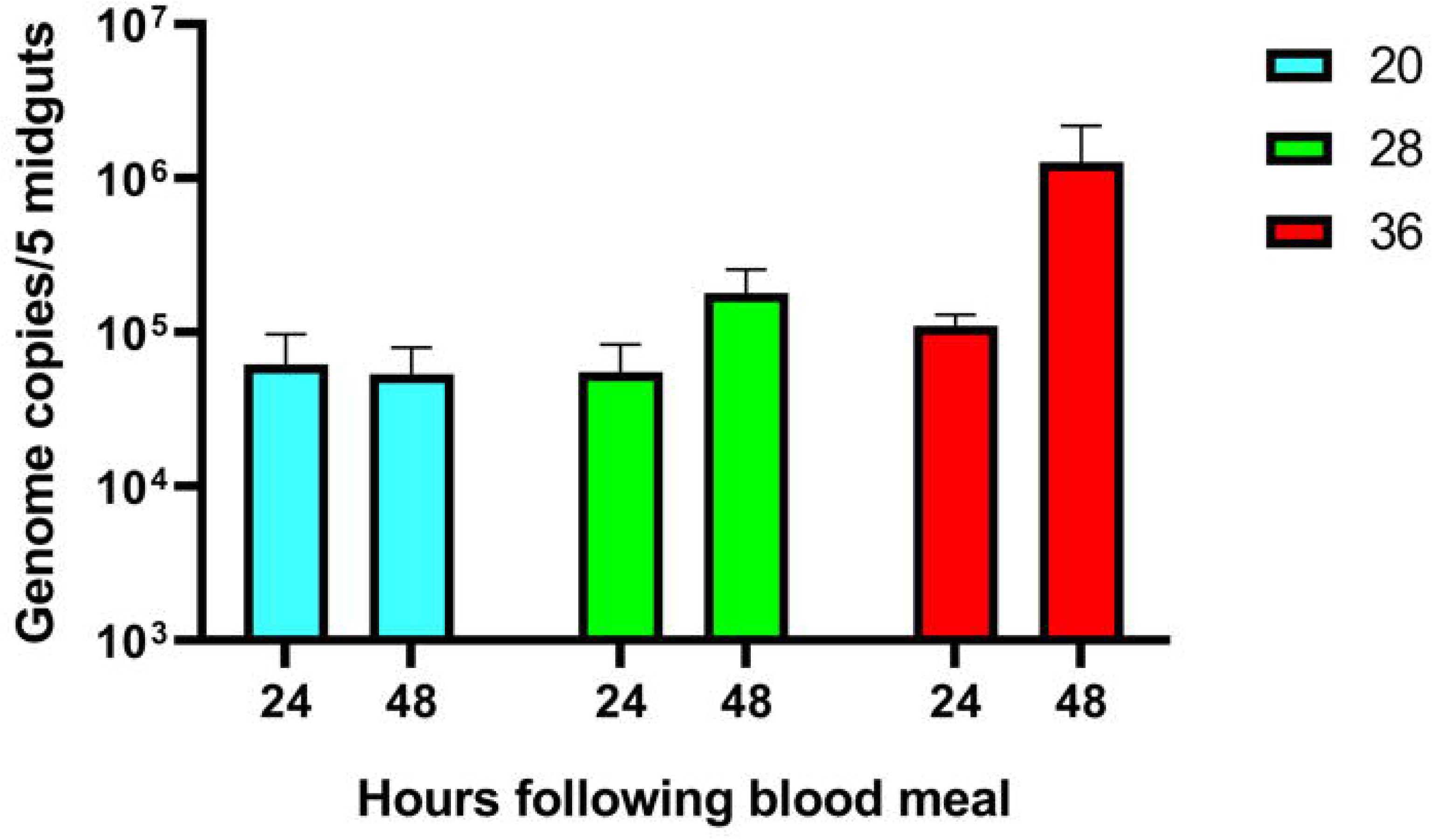
Quantification of ZIKV RNA in infected mosquitoes housed at 20, 28, and 36°C at two 24 and 48hs times. The average number of ZIKV genome copies present in pools of 5 midgut samples and standard error are shown.

### The effect of temperature on gene expression in unexposed mosquitoes

Gene expression patterns were influenced by both temperature and time post-blood feed. In general, the principal-component analysis (PCA) plot showed a high degree of reproducibility among the replicate samples within each temperature treatment. At 24 hpf, temperature clearly distinguished gene expression profiles (Supplementary Figure S1 A). However, by 48 hpf, mosquitoes housed at 28°C and 36°C had more similar gene expression profiles relative to mosquitoes housed at 20°C (Figure 2A). The vast majority of genes expressed in the midgut were similar from mosquitoes housed across the three temperature treatments (10,459 at 24 hpf, Supplementary Figure S1 B; 10,850 at 48 hpf, Figure 2B). Euclidean distance heatmap analysis demonstrate that the differentially expressed genes at 28°C and 36°C cluster more closely with one another than with the 20°C samples at 48 hpf (Figure 2C). We identified 1665 differentially expressed genes (q-value < 0.05) between mosquitoes held at 20°C and 28°C for 24 hr and 3634 by 48 hr (Data Sheet 1). In contrast, when comparing mosquitoes held at 36°C and 28°C, we identified 1697 and 1695 differentially expressed genes after 24 hr and 48 hr, respectively (Data Sheet 2). In order to produce a more stringent analysis, we produced a volcano plot indicating a total of 70 and 297 upregulated genes (log_2_(Fold Change) > 2 and q-value < 0.05) in mosquitoes housed at 20°C for 24 hr and 48 hr, respectively, compared to 28°C. For downregulated genes (log_2_(Fold Change) < −2 and q-value < 0.05), we found 53 and 317 after 24 hr and 48 hr, respectively (Supplementary Figure S 1D and Figure 2D). We did not find as strong of an effect of exposure time when mosquitoes were housed at 36°C, with only 19 and 39 genes up-regulated and 89 and 79 genes down-regulated after 24 and 48 hr, respectively (Supplemental Figure S 1D and Figure 2D).

**Figure 2.**
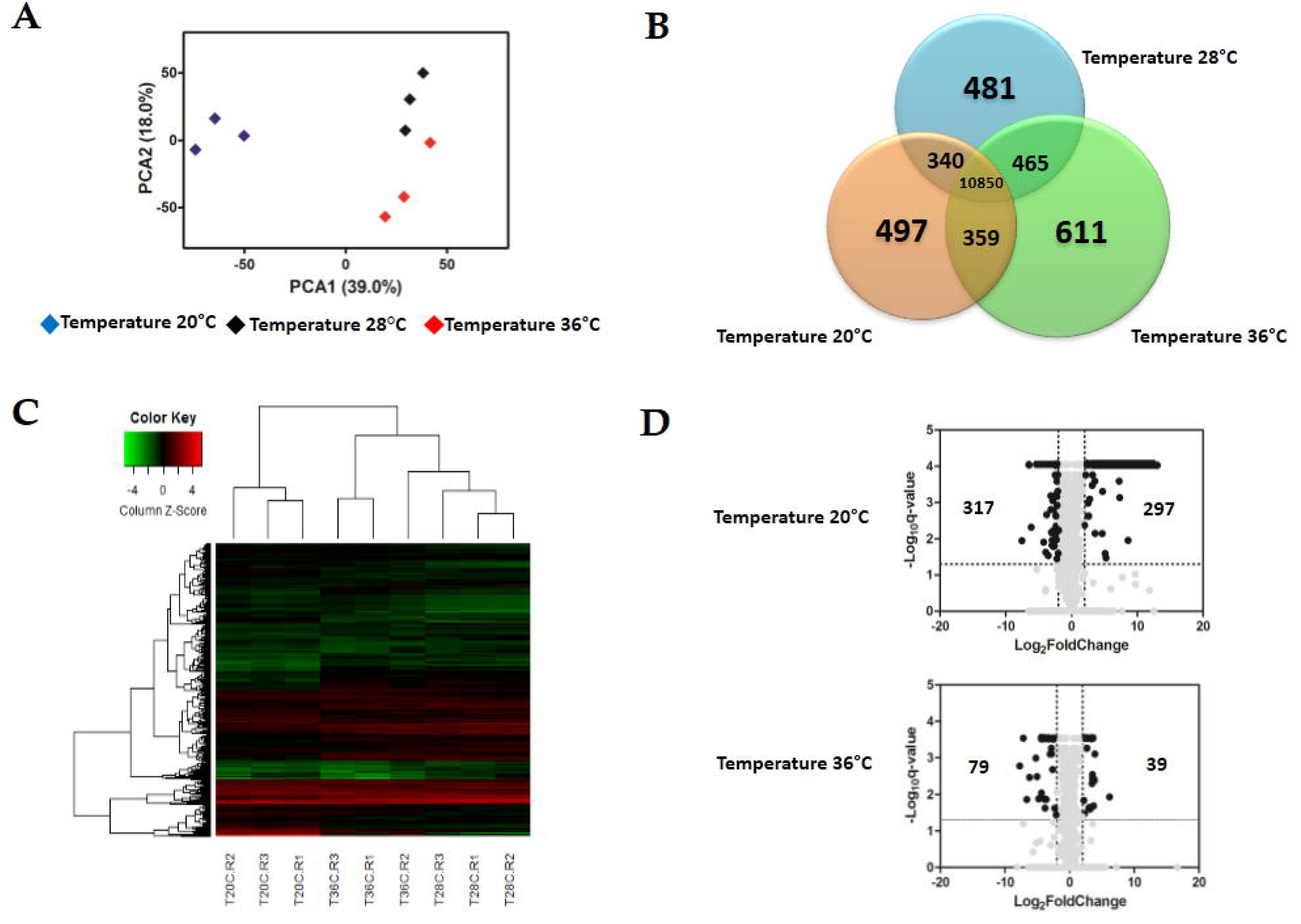
Effect of temperature in the expression profile of uninfected blood fed *Ae. aegypti*. **(A)** Principal Component Analysis (PCA) plot showing the global gene expression profiles **(B)** Heatmap plot showing local differences (TC20R1: represents the replicate 1 for the temperature 20°C) **(C)** Venn Diagram reporting the number of specific and shared genes **(D)** Volcano plot representing the differential gene expression from RNAseq samples from *Aedes aegypti* exposed to three different constant temperatures (20°C, 28°C, 36°C) at 48 hrs.

To further define the effect of temperature on mosquito cellular and physiological responses, we sorted the top 20 up- or down-regulated genes at both the cool and hot temperatures using mosquitoes maintained at 28°C as our standard (Table 1). The cool temperature produced a larger response, up-regulating genes hundreds of times compared to standard conditions, whereas the 36°C treatment had a more moderate effect. The transcript with the highest enrichment at 20°C was protein-G12 (1459-fold), whereas, a serine protease easter was the most down-regulated (187-fold). For mosquitoes housed at 36°C, heat shock protein 70 (HSP 70) was the transcript most enriched (71-fold) relative to 28°C, while facilitated trehalose transporter Tret1 was the most down-regulated gene (212-fold). Surprisingly, both cool and warm temperature treatments induced some of the same genes to be differentially regulated. Six genes (protein G12, serine protease SP24D, and chymotrypsin-2) were up regulated in mosquitoes housed at both 20°C and 36°C when compared to those at 28°C, while six genes (serine protease easter, facilitated trehalose transporter Tret1, venom protease, solute carrier family 2 facilitated glucose transporter member 3, and tyrptase) were down regulated.

**Table 1.** Top 20 up and down-regulated genes of uninfected Ae. aegypti in response to low (20°C) and to high (36°C) temperature for 48 hr post-feeding blood (relative to standard rearing temperature of 28°C).

| GENES UP REGULATED |  | TEMPERATURE 20°C |  | TEMPERATURE 36°C |  |
| --- | --- | --- | --- | --- | --- |
| Gene ID | Gene Description | FoldChange | q-value | FoldChange | q-value |
| XP_021701760.1 | Protein G12 | 1459.541989 | 0.0000886602 | 11.11657146 | 0.000290131 |
| XP_021698904.1 | Chymotrypsin-2 | 857.284352 | 0.0000886602 |  |  |
| XP_001660827.1 | Protein G12 | 603.3619885 | 0.0000886602 | 7.4143488679 | 0.000290131 |
| XP_001650490.2 | Alpha-N-acetylgalactosaminidase | 548.7746389 | 0.0000886602 |  |  |
| XP_001659962.1 | Serine protease SP24D | 473.2635493 | 0.0000886602 | 11.29286782 | 0.000290131 |
| XP_021712126.1 | Protein G12-like | 442.0421098 | 0.0000886602 |  |  |
| XP_021701761.1 | Protein G12 | 402.0329675 | 0.0000886602 | 6.3523041907 | 0.000548702 |
| XP_001663002.1 | Trypsin 3A1-like | 384.5293743 | 0.0110252 |  |  |
| XP_021705369.1 | Beta-galactosidase | 315.6714553 | 0.0000886602 |  |  |
| XP_001656375.1 | Protein G12 | 245.4048583 | 0.0000886602 |  |  |
| XP_001656377.1 | Protein G12 isoform X2 | 221.8114454 | 0.0000886602 | 5.0806039301 | 0.000290131 |
| XP_021703511.1 | Lysosomal alpha-mannosidase isoform X2 | 217.023882 | 0.0000886602 |  |  |
| XP_001647937.2 | Phosphoenolpyruvate carboxykinase [GTP] | 180.2043466 | 0.0000886602 |  |  |
| XP_001659796.1 | Beta-1,3-glucan-binding protein | 159.2092116 | 0.000730421 |  |  |
| XP_021712858.1 | Protein G12 isoform X2 | 149.8743826 | 0.000256108 |  |  |
| XP_001657509.1 | Vitellogenin-A1-like | 138.4047839 | 0.0000886602 |  |  |
| XP_001659961.1 | Chymotrypsin-2 | 110.7367288 | 0.0000886602 | 8.596677982 | 0.000290131 |
| XP_001652194.1 | Protein singed wings 2 | 84.66536466 | 0.0000886602 |  |  |
| XP_001660818.2 | Vitellogenin-A1 | 67.7965212 | 0.0000886602 |  |  |
| XP_001654186.1 | Glutamine synthetase 1 mitochondrial | 65.69082595 | 0.0000886602 |  |  |
| XP_021693649.1 | Heat shock protein 70 A1 |  |  | 71.53314665 | 0.0118095 |
| XP_001660673.2 | Trypsin 5G1-like |  |  | 14.98866514 | 0.0007852 |
| XP_001663776.1 | Protein G12 |  |  | 14.35856831 | 0.00416417 |
| XP_001658359.2 | Brachyurin |  |  | 12.24869336 | 0.000290131 |
| XP_021701762.1 | Protein G12-like |  |  | 12.07440339 | 0.00379042 |
| XP_001663895.2 | Trypsin 5G1 |  |  | 11.04851064 | 0.00505811 |
| XP_001658471.2 | Mite group 2 allergen Gly |  |  | 10.38200486 | 0.000290131 |
| XP_011493274.2 | Extensin |  |  | 9.222769233 | 0.000290131 |
| XP_001659492.2 | Serine protease SP24D |  |  | 7.5082672369 | 0.000290131 |
| XP_001661186.2 | Protein G2 |  |  | 6.9220050786 | 0.000290131 |
| XP_001651623.2 | Surface antigen CRP170 |  |  | 6.151678044 | 0.000290131 |
| XP_021712126.1 | Protein G12-like |  |  | 5.6022641153 | 0.000548702 |
| XP_001660908.1 | Maltase 1 |  |  | 5.3506716535 | 0.000290131 |
| XP_011492940.1 | Peptidoglycan-recognition protein SC-2 |  |  | 5.030075309 | 0.000290131 |

| GENES DOWN REGULATED |  | TEMPERATURE 20°C |  | TEMPERATURE 36°C |  |
| --- | --- | --- | --- | --- | --- |
| Gene ID | Gene Description | FoldChange | q-value | FoldChange | q-value |
| XP_001652078.2 | Serine protease easter | 187.8360285 | 0.0112238 | 33.06859855 | 0.0032544 |
| XP_001658147.1 | Facilitated trehalose transporter Tret1 | 39.72595509 | 0.0000886602 | 212.5163457 | 0.00166701 |
| XP_001652075.2 | Venom protease | 31.87315659 | 0.0000886602 | 21.42212957 | 0.000290131 |
| XP_001663064.2 | UDP-glucuronosyltransferase 2B18 | 28.70798168 | 0.0000886602 |  |  |
| XP_001652079.1 | Serine protease easter | 18.68203877 | 0.0124204 | 15.08894466 | 0.0134155 |
| XP_001655069.1 | Queuine tRNA-ribosyltransferase accessory subunit 2 | 15.34875785 | 0.0000886602 |  |  |
| XP_001656516.2 | Xaa-Pro aminopeptidase ApepP | 15.23840501 | 0.0000886602 |  |  |
| XP_001649987.2 | Synaptic vesicle glycoprotein 2B | 13.81630566 | 0.00215644 |  |  |
| XP_021706766.1 | Glycerol-3-phosphate dehydrogenase mitochondrial isoform X1 | 13.39803945 | 0.0000886602 |  |  |
| XP_021699787.1 | Solute carrier family 2 facilitated glucose transporter member 3 | 13.34678167 | 0.0000886602 | 99.27028299 | 0.0138613 |
| XP_001655305.2 | Protein MAK16 homolog A | 12.43254717 | 0.0000886602 |  |  |
| XP_001649768.1 | Exosome complex component RRP43 | 11.98593043 | 0.0000886602 |  |  |
| XP_001653029.1 | Protein takeout | 11.66609644 | 0.0000886602 |  |  |
| XP_021698953.1 | CAD protein | 11.58599466 | 0.0291848 |  |  |
| XP_001656680.1 | NHP2-like protein 1 homolog | 10.79948043 | 0.0000886602 |  |  |
| XP_001655729.2 | Tryptase | 10.69882162 | 0.0000886602 | 13.05191869 | 0.000290131 |
| XP_001662512.1 | Nucleoside diphosphate kinase | 10.5849313 | 0.0000886602 |  |  |
| XP_001663497.1 | Protein lethal(2) essential for life | 10.23543137 | 0.0000886602 |  |  |
| XP_001655105.1 | Sorbitol dehydrogenase | 10.21211653 | 0.0000886602 |  |  |
| XP_021699336.1 | Nucleoside diphosphate kinase-like | 10.07585248 | 0.0000886602 |  |  |
| XP_001652056.2 | Vitellogenic carboxypeptidase |  |  | 37.57757336 | 0.00101347 |
| XP_001655104.1 | Sorbitol dehydrogenase |  |  | 27.67324652 | 0.013266 |
| XP_001663173.2 | Solute carrier family 22 member 8 |  |  | 19.16072174 | 0.000290131 |
| XP_021702456.1 | Sodium-coupled monocarboxylate transporter 1 isoform X1 |  |  | 18.47599301 | 0.000290131 |
| XP_001651077.2 | Synaptic vesicle glycoprotein 2B |  |  | 17.7140231 | 0.000290131 |
| XP_001656519.2 | Solute carrier family 22 member 21 isoform X2 |  |  | 14.01519585 | 0.000290131 |
| XP_001662495.2 | Lipase 1 |  |  | 13.73780875 | 0.000290131 |
| XP_021706833.1 | Sodium-coupled monocarboxylate transporter 1 isoform X2 |  |  | 13.50049126 | 0.000290131 |
| XP_021698236.1 | Solute carrier organic anion transporter family member 2A1 |  |  | 11.93072658 | 0.000290131 |
| XP_021702682.1 | Acidic amino acid decarboxylase GADL1 isoform X3 |  |  | 11.91023521 | 0.000290131 |
| XP_001649855.2 | Sodium/potassium/calcium exchanger 4 |  |  | 11.67450924 | 0.000290131 |
| XP_021710339.1 | Synaptic vesicle glycoprotein 2C |  |  | 10.26811889 | 0.000290131 |
| XP_001662720.1 | Putative transporter SVOPL |  |  | 10.08241963 | 0.000290131 |
| XP_021713275.1 | Synaptic vesicle glycoprotein 2A |  |  | 8.744949179 | 0.000290131 |

All differentially expressed genes in the midgut at 48 hpf were submitted for gene ontology (GO) analysis to identify cellular processes that were most perturbed in response to temperature treatment as revealed by the transcriptome profile. GO analysis of the enriched and depleted transcripts from mosquitoes housed at 20°C revealed that 20 enriched GO terms were related to oxidation-reduction processes and 111 depleted GO terms were involved in gene expression, RNA processing, metabolic processes and generation of energy (Figure 3). Mosquitoes housed at 36°C displayed up-regulated expression related to amine metabolism and cell redox homeostasis processes and down-regulated expression of genes associated with metabolic processes, cellular respiration, and energy derivation by oxidation of organic compounds (Figure 3).

**Figure 3.**
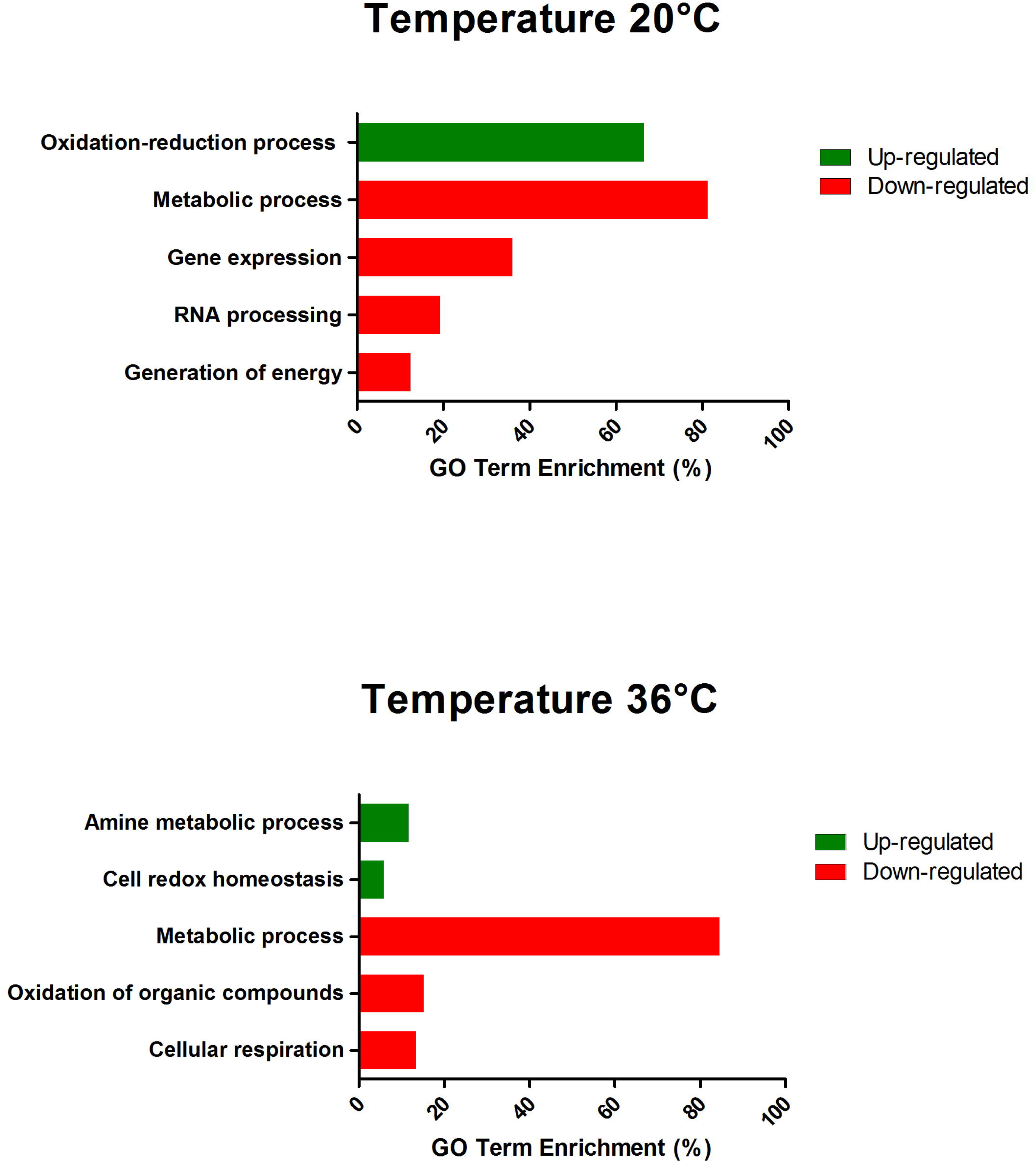
GO term enrichment analysis of differentially expressed genes in response to exposed of Zika-infected *A. aegypti* in the lowest (20°C) and in the highest (36°C) temperature in the category of biological process for up and down-regulated genes at 48 hs.

### The effect of temperature on gene expression in ZIKV-exposed mosquitoes

The gene expression profiles in ZIKV-exposed mosquitoes were also significantly affected by environmental temperature and time post-blood feed. In general, the effect of temperature on differential gene expression was similar to patterns observed in non-ZIKV exposed blood-fed control mosquitoes outlined above. For example, PCA and heatmap analyses on differential gene expression from mosquito midguts at 24 hpf illustrated three distinct groups separating by temperature treatment (Supplementary Figure S2 A and S2 C). At 48 hpf, gene expression in mosquitoes housed at 20°C was more distinct than those housed at 28°C and 36°C (Figure 4A and 4C). Further, Venn diagrams demonstrate that 1416 and 10,786 genes were expressed across all temperatures at 24 hpf (Supplementary Figure S2 B) and 48 hpf (Figure 4B), respectively, with the highest overlap in gene expression occurring in mosquitoes housed at 28°C and 36°C at both time points (Supplementary Figure S2 B, Figure 4B). Finally, as seen in the absence of ZIKV infection, at 24 hpf, only 1669 and 1797 genes were differentially expressed in mosquitoes housed at 20°C and 36°C, respectively, relative to those housed at 28°C. By 48 hpf, we observed a general increase in the number of genes differentially expressed at 20°C (3056, Data Sheet 3) and a general decrease at 36°C (1518, Data Sheet 4) relative to mosquitoes housed at 28°C. When we evaluated differential expression through a volcano plot, the profile was similar to mosquitoes that were not exposed to ZIKV (Supplemental Figure S 2D and Figure 4D).

**Figure 4.**
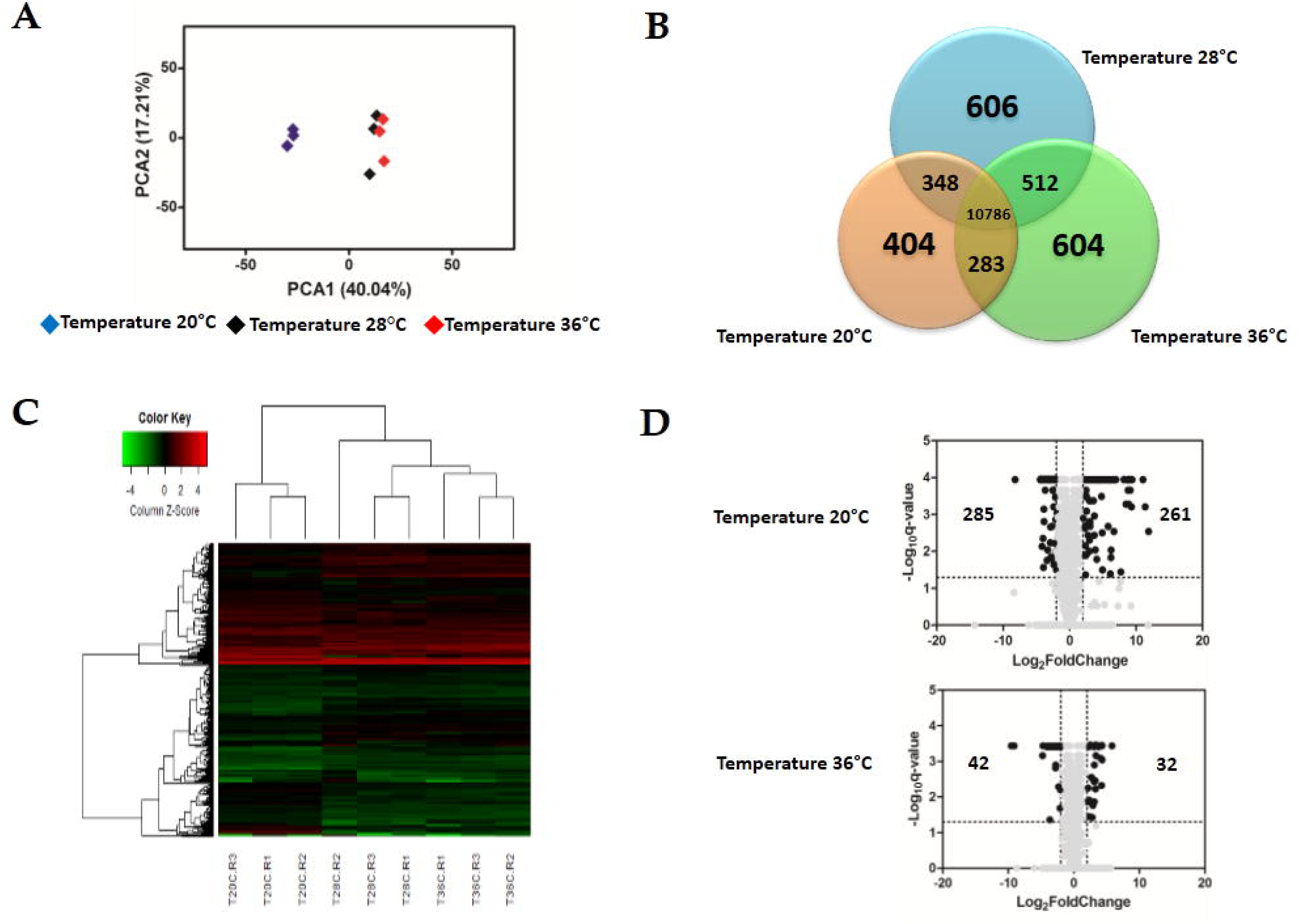
Effect of the temperature in the expression profile of Zika-exposed *A. aegypti*. **(A)** Principal Component Analysis (PCA) plot showing the global gene expression profiles **(B)** Heatmap plot showing local differences (TC20R1: represents the replicate 1 for the temperature 20°C) **(C)** Venn Diagram reporting the number of specific and shared genes **(D)** Volcano plot representing the differential gene expression from RNAseq samples from infected *Aedes aegypti* exposed in three different constant temperatures (20°C, 28°C, 36°C) at 48 hs.

Several genes that strongly increased at 20°C remained among the top 20 differentially expressed after ZIKV exposure. Lysosomal alpha-mannosidase (XP_021703511.1), two vitellogenins (XP_001660818.2, XP_001657509.1), phosphoenolpyruvate carboxykinase (XP_001647937.2), proteins G12 (XP_021712126.1, XP_021701760.1, XP_001660827.1, XP_021701761.1, XP_001656377.1, XP_001656375.1), beta-galactosidase (XP_021705369.1), alpha-N-acetylgalactosaminidase (XP_001650490.2), serine protease SP24D (XP_001659962.1) and chymotrypsin-2 (XP_021698904.1) are some of genes that remained among 20-top changed genes by cold temperature in the ZIKV-infection condition. We also saw that although glutamine synthetase (XP_001654186.1) and trypsin (XP_001663002.1) were not listed among 20 most differentially expressed, they showed enrichment of 50 and 74-fold, respectively. Further, facilitated trehalose transporter Tret1, found to be the most down-regulated in unexposed mosquitoes kept at 36°C, was also the most down-regulated (25-fold) in ZIKV-exposed mosquitoes at the same temperature. Finally, serine protease SP24D (XP_001659962.1) and some G12 proteins (XP_021701760.1, XP_021701761.1, XP_001660827.1, XP_001656377.1) that were enriched at both cool and warm temperatures, remained enriched at these temperatures in ZIKV exposed mosquitoes (Table 2).

**Table 2.** Top 20 up and down-regulated genes of ZIKV-exposed *Ae. aegypti* in response to low (20°C) and to high (36°C) temperature for 48 hs post-feeding blood relative to standard insectary conditions (28°C).

| GENES UP REGULATED |  | TEMPERATURE 20°C |  | TEMPERATURE 36°C |  |
| --- | --- | --- | --- | --- | --- |
| Gene ID | Gene Description | FoldChange | q-value | FoldChange | q-value |
| XP_001657509.1 | Vitellogenin-A1-like | 3748.76585 | 0.00288439 |  |  |
| XP_001660818.2 | Vitellogenin-A1 | 2644.35472 | 0.000624083 |  |  |
| XP_021701760.1 | Protein G12 | 2081.92156 | 0.000113659 | 18.9560964 | 0.00036736 |
| XP_001660827.1 | Protein G12 | 668.331471 | 0.000113659 | 9.8533206 | 0.00036736 |
| XP_021701762.1 | Protein G12-like | 626.881959 | 0.000624083 | 9.79462325 | 0.00599791 |
| XP_021712126.1 | Protein G12-like | 580.732514 | 0.000113659 | 8.68875819 | 0.00126501 |
| XP_001663776.1 | Protein G12 | 519.780968 | 0.000113659 | 10.3753637 | 0.000690526 |
| XP_021701761.1 | Protein G12 | 498.942585 | 0.000113659 | 9.76777521 | 0.00036736 |
| XP_021698904.1 | Chymotrypsin-2 | 476.004037 | 0.000113659 |  |  |
| XP_001657506.2 | Vitellogenin-A1-like | 467.414928 | 0.000526591 |  |  |
| XP_001656377.1 | Protein G12 isoform X2 | 400.561513 | 0.000113659 | 8.94238711 | 0.00036736 |
| XP_021702099.1 | Probable nuclear hormone receptor HR3 isoform X5 | 356.184271 | 0.000526591 |  |  |
| XP_001656375.1 | Protein G12 | 315.925373 | 0.000113659 |  |  |
| XP_001659962.1 | Serine protease SP24D | 315.50958 | 0.000113659 | 8.63508273 | 0.00036736 |
| XP_021705369.1 | Beta-galactosidase | 310.614205 | 0.000113659 |  |  |
| XP_001650490.2 | Alpha-N-acetylgalactosaminidase | 292.584595 | 0.000113659 |  |  |
| XP_001652055.1 | Vitellogenic carboxypeptidase-like | 206.875645 | 0.0359751 |  |  |
| XP_001652056.2 | Vitellogenic carboxypeptidase | 123.386436 | 0.000113659 |  |  |
| XP_001647937.2 | Phosphoenolpyruvate carboxykinase [GTP] | 106.495599 | 0.000113659 |  |  |
| XP_021703511.1 | Lysosomal alpha-mannosidase isoform X2 | 103.072096 | 0.000113659 |  |  |
| XP_001652358.2 | Peritrophin-1 |  |  | 55.8211523 | 0.00036736 |
| XP_021697715.1 | Peritrophin-1-like |  |  | 16.3831402 | 0.00036736 |
| XP_001658471.2 | Mite group 2 allergen Gly d 2.01 |  |  | 15.940003 | 0.00036736 |
| XP_001663895.2 | Trypsin 5G1 |  |  | 9.16859073 | 0.00365688 |
| XP_001660673.2 | Trypsin 5G1-like |  |  | 8.69411995 | 0.0136618 |
| XP_001661186.2 | Protein G12 |  |  | 7.2863687 | 0.00036736 |
| XP_001663102.2 | Malate synthase |  |  | 6.86576273 | 0.00036736 |
| XP_001663439.2 | Collagenase |  |  | 6.39130263 | 0.00279659 |
| XP_021707618.1 | Probable chitinase 2 |  |  | 6.27684681 | 0.00036736 |
| XP_001651623.2 | Surface antigen CRP170 |  |  | 5.38042449 | 0.00036736 |
| XP_001653091.2 | Trypsin alpha-3 |  |  | 5.26940475 | 0.00036736 |
| XP_001659961.1 | Chymotrypsin-2 |  |  | 5.00475692 | 0.00036736 |

| GENES DOWN REGULATED |  | TEMPERATURE 20°C |  | TEMPERATURE 36°C |  |
| --- | --- | --- | --- | --- | --- |
| Gene ID | Gene Description | FoldChange | q-value | FoldChange | q-value |
| XP_001663064.2 | UDP-glucuronosyltransferase 2B18 | 22.0442425 | 0.000113659 | 5.62311675 | 0.00036736 |
| XP_011493503.2 | H/ACA ribonucleoprotein complex subunit 1 | 20.5197322 | 0.000113659 |  |  |
| XP_001654398.2 | rRNA 2'-O-methyltransferase fibrillar | 19.1980783 | 0.000113659 |  |  |
| XP_001659197.1 | DNA-directed RNA polymerase III subunit RPC10 | 18.0109204 | 0.00732014 |  |  |
| XP_001658147.1 | Facilitated trehalose transporter Tret1 | 17.354405 | 0.000113659 | 25.7984056 | 0.00036736 |
| XP_001663005.2 | Periodic tryptophan protein 1 homolog | 16.363733 | 0.000113659 |  |  |
| XP_021698953.1 | CAD protein | 16.0097625 | 0.00443189 |  |  |
| XP_001655305.2 | Protein MAK16 homolog A | 15.3024421 | 0.000113659 |  |  |
| XP_001656228.2 | COX assembly mitochondrial protein 2 homolog | 14.7057657 | 0.000721477 |  |  |
| XP_021706766.1 | Glycerol-3-phosphate dehydrogenase mitochondrial isoform X1 | 14.6844772 | 0.000113659 |  |  |
| XP_001649209.1 | RRP15-like protein isoform X2 | 14.5214136 | 0.00156177 |  |  |
| XP_001652303.1 | mRNA turnover protein 4 homolog | 14.3570755 | 0.000113659 |  |  |
| XP_001662723.2 | Titin homolog | 14.3017521 | 0.000113659 |  |  |
| XP_021699084.1 | Proton-coupled amino acid transporter 1-like | 14.2666035 | 0.000113659 |  |  |
| XP_001658562.1 | Activator of basal transcription 1 | 13.971742 | 0.000113659 |  |  |
| XP_021695012.1 | 46 kDa FK506-binding nuclear protein | 13.8346093 | 0.000113659 |  |  |
| XP_001660583.2 | Glutamate-rich WD repeat-containing protein 1 | 13.2594589 | 0.000113659 |  |  |
| XP_021703839.1 | Protein Notchless | 13.0245353 | 0.000113659 |  |  |
| XP_001656680.1 | NHP2-like protein 1 homolog | 12.0716418 | 0.000113659 |  |  |
| XP_001655996.2 | H/ACA ribonucleoprotein complex non-core subunit NAF1 | 11.9572191 | 0.000113659 |  |  |
| XP_001662495.2 | Lipase 1 |  |  | 16.3418567 | 0.00036736 |
| XP_021698905.1 | Chymotrypsin-2 |  |  | 13.9310298 | 0.00036736 |
| XP_001649987.2 | Synaptic vesicle glycoprotein 2B |  |  | 12.6667105 | 0.0434966 |
| XP_021702682.1 | Acidic amino acid decarboxylase GADL1 isoform X3 |  |  | 11.6866538 | 0.00036736 |
| XP_001649098.2 | Probable cytochrome P450 9f2 |  |  | 8.68297843 | 0.00036736 |
| XP_021699787.1 | Solute carrier family 2 facilitated glucose transporter member 3 |  |  | 7.85777033 | 0.00036736 |
| XP_001652075.2 | Venom protease |  |  | 6.9627634 | 0.00153067 |
| XP_021702456.1 | Sodium-coupled monocarboxylate transporter 1 isoform X1 |  |  | 6.31621377 | 0.00036736 |
| XP_001651954.2 | Trypsin |  |  | 6.12067175 | 0.00036736 |
| XP_001649855.2 | Sodium/potassium/calcium exchanger 4 |  |  | 5.8573577 | 0.00036736 |
| XP_021704942.1 | Excitatory amino acid transporter 1 |  |  | 5.56974007 | 0.00036736 |
| XP_021708608.1 | Putative helicase MOV-10 |  |  | 5.4055447 | 0.00036736 |
| XP_001656046.1 | Alanine--glyoxylate aminotransferase 2-like |  |  | 5.35883724 | 0.00036736 |
| XP_021698609.1 | Chymotrypsin-2 |  |  | 5.27740974 | 0.00036736 |
| XP_001661015.2 | Putative alpha-L-fucosidase |  |  | 5.02108797 | 0.00036736 |
| XP_001658000.3 | Glutathione S-transferase 1 |  |  | 5.0085396 | 0.00517339 |
| XP_021704288.1 | Phosphotriesterase-related protein |  |  | 4.96319797 | 0.00036736 |
| XP_001661388.1 | Chymotrypsin-1 |  |  | 4.7758913 | 0.00036736 |

Despite these similarities, we did note some key differences between the top 20 genes most differentially expressed in unexposed (Table 1) and ZIKV-exposed mosquitoes (Table 2), with the greatest change in expression reflected in mosquitoes housed at 20°C and 36°C at 48 hr relative to those housed at 28°C. In unexposed mosquitoes at 20°C vitellogenin-A1-like (XP_001657509.1) and vitellogenin-A1 (XP_001660818.2) were up-regulated, yet ZIKV exposure amplified the enrichment from 138 and 68 fold to 3748 and 2644 fold (Table 1 and 2). Surprisingly in ZIKV exposed mosquitoes we did not detect a large up-regulation of Hsp70 at 36°C like we did in the unexposed population.

The GO analysis of differentially expressed genes at 48 hpf in ZIKV-exposed mosquitoes demonstrated distinct effects of cool and warm temperatures on cellular and metabolic function relative to mosquitoes housed at 28°C (Figure 5). Further, the functions of these differentially expressed genes were in part different from unexposed mosquitoes housed at these temperatures (Figure 3). For example,, oxidative-reduction processes maintained at 20°C were no longer enriched as seen in unexposed mosquitoes. Instead, ZIKV-exposed mosquitoes housed at 20°C had significant enrichment of the Toll signaling pathway, a known anti-dengue pathway in *Ae. aegypti* (Tchankouo-Nguetcheu et al., 2010). Additionally, endosome transport, Ras protein signal transduction, actin cytoskeleton organization, epithelial tube morphogenesis, pH reduction, and proteolysis were enriched. In contrast nuclear transport, regulation of viral reproduction, “*de novo*” protein folding, generation of energy, gene expression, RNA processing, and metabolic processes were down-regulated in addition to expression associated with gene expression, RNA processing, and metabolic processes observed in unexposed counterparts at this temperature (Figure 5). ZIKV-exposed mosquitoes housed at 36°C no longer had significant enrichment in genes associated with cellular amine processes and had significant depletion in genes associated with hexose and phosphate metabolic processes, as well as the generation of precursor metabolites and energy (Figure 5).

**Figure 5.**
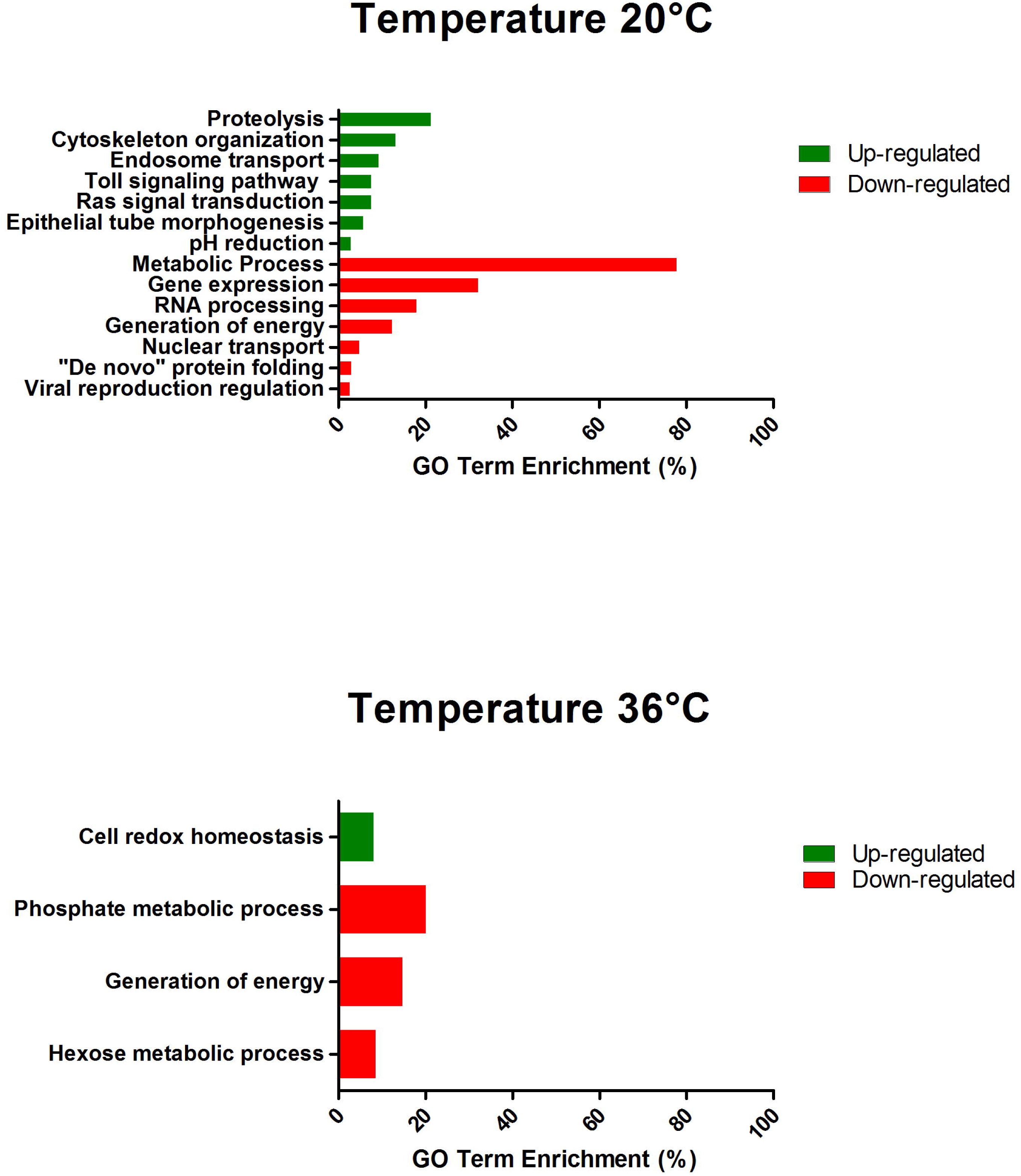
GO term enrichment analysis of differentially expressed genes in response to exposed of Zika-infected *A. aegypti* in the lowest (20°C) and in the highest (36°C) temperature in the category of biological process for up and down-regulated genes at 48 hs.

### Effects of temperature on oxidative stress and innate immune mechanisms

To further explore the effects of temperature on uninfected and ZIKV-exposed mosquitoes, we highlight those involved in managing oxidative stress, innate immunity, and apoptosis at both 20°C and 36°C compared to 28°C at 48 hpf (Data Sheet 5 and 6, Supplementary Figure S3 and S4). Virus infection is modified by complex responses related to detoxification of the blood meal, metabolism, immunity, and apoptosis in some systems. (Sanders et al., 2005; Girard et al., 2010; Tchankouo-Nguetcheu et al., 2010; Colpitts et al., 2011; Wang et al., 2012; Neill et al., 2015; Eng et al., 2016). Further, these responses need not respond equivalently to temperature variation. The majority of genes involved in managing oxidative stress, innate immunity, and apoptosis exhibited qualitatively different patterns in gene expression in response to cool and warm temperatures in uninfected and ZIKV-exposed mosquitoes. However, while significant, the majority of these differences were very subtle (< 2.0 fold; Supplementary Figure S3 and S4). We did observe components of the melanization and Toll pathways to be modestly expressed (>2.0 fold; Supplementary Figure S3 and S4) in response to temperature. An isoform of phenoloxidase (XP_021699380.1), a c-type lectin (XP_001661643.1), and the Toll receptor 6 (TLR6) (XP_021712805.1) were significantly enriched in both uninfected and ZIKV-exposed mosquitoes housed at 20°C.

### Temperature has subtle effects on mosquito response to ZIKV infection

To investigate the effects of ZIKV-exposure on global gene expression patterns from mosquito midguts early on in the infection process, we compared gene expression between ZIKV-exposed and unexposed mosquitoes within each temperature treatment. PCA plots demonstrate that ZIKV exposure does not significantly alter mosquito transcription at 24 and 48 hpf (Supplementary Figure S5, Figure 6) within a given temperature treatment. However, the overall number of differentially expressed genes between ZIKV-exposed and unexposed mosquitoes varied across temperature treatments. For example, at 24 hpf, we observed a total of 225, 154, and 161 genes differentially expressed at 20°C, 28°C, and 36°C, respectively, between ZIKV-exposed and unexposed mosquitoes (Data Sheet 7, 8 and 9). We identified only two proteins - a sodium/potassium/calcium exchanger 4 (XP_001649855.2) and an uncharacterized protein (XP_001654261.2) - that were up-regulated in ZIKV-exposed mosquitoes at all temperatures (Supplementary Figure S6 A) and one protein - chymotrypsin-2 (XP_021698609.1) that was down-regulated (Supplementary Figure S6 B). At 48 hpf, we observed more genes to be differentially expressed (1188 genes) in mosquitoes housed at 20°C between ZIKV-exposed and unexposed mosquitoes, and only 180 and 50 genes at 28°C and 36°C, respectively (Data Sheet 7, 8 and 9). Only one uncharacterized protein (XP_001658660.2) was up-regulated in the ZIKV-exposed mosquitoes at all temperatures (Figure 7A) at this sampling time point. These results indicate that while the physiological responses of mosquito midguts to ZIKV exposure early in the infection process are similar within a given temperature treatment, these responses are significantly distinct across different environmental temperatures.

**Figure 6.**
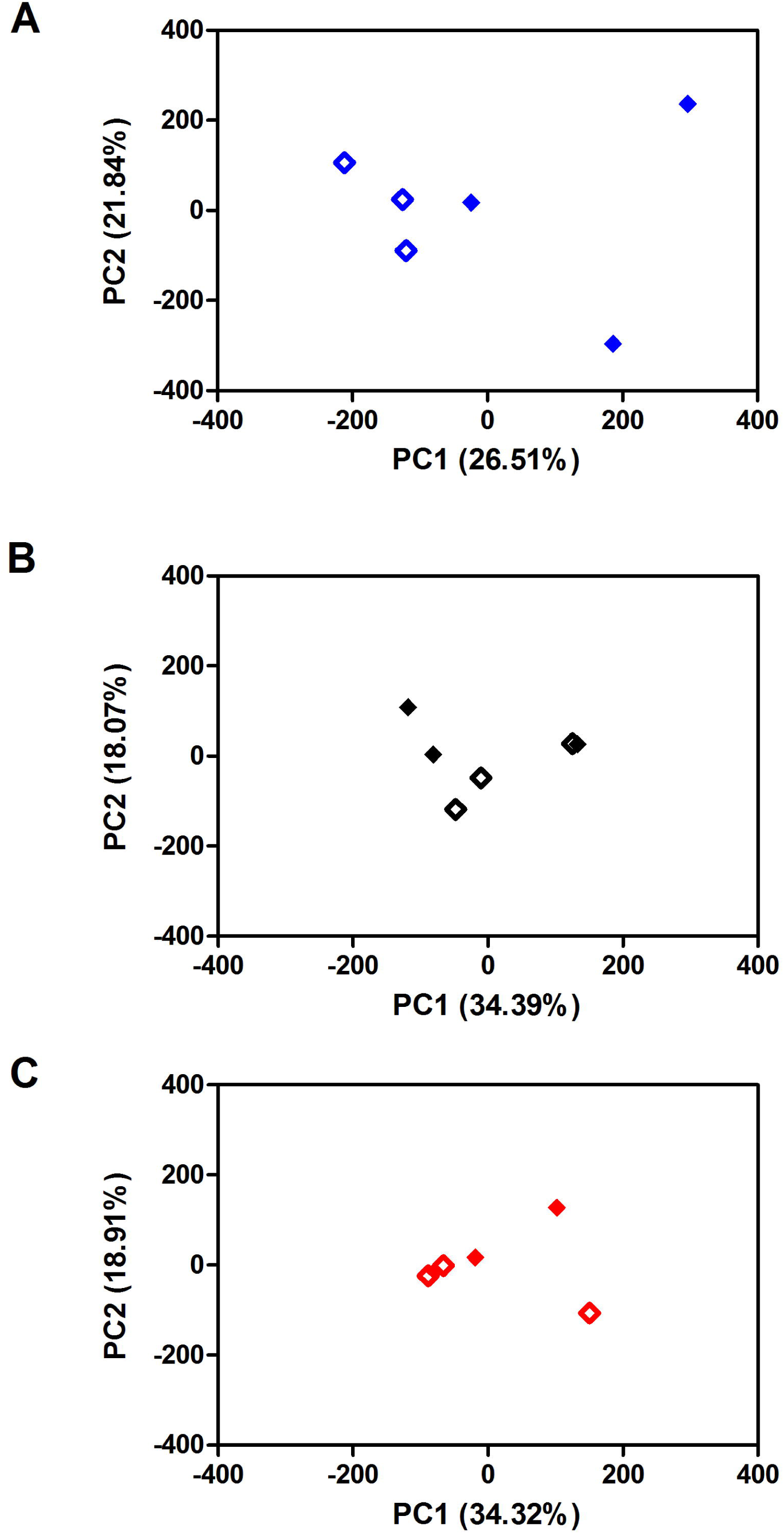
The Principal Component Analysis (PCA) of the effect of the Zika exposure in three different constant temperatures (A) 20°C (B) 28°C (C) 36°C at 48hs.

**Figure 7.**
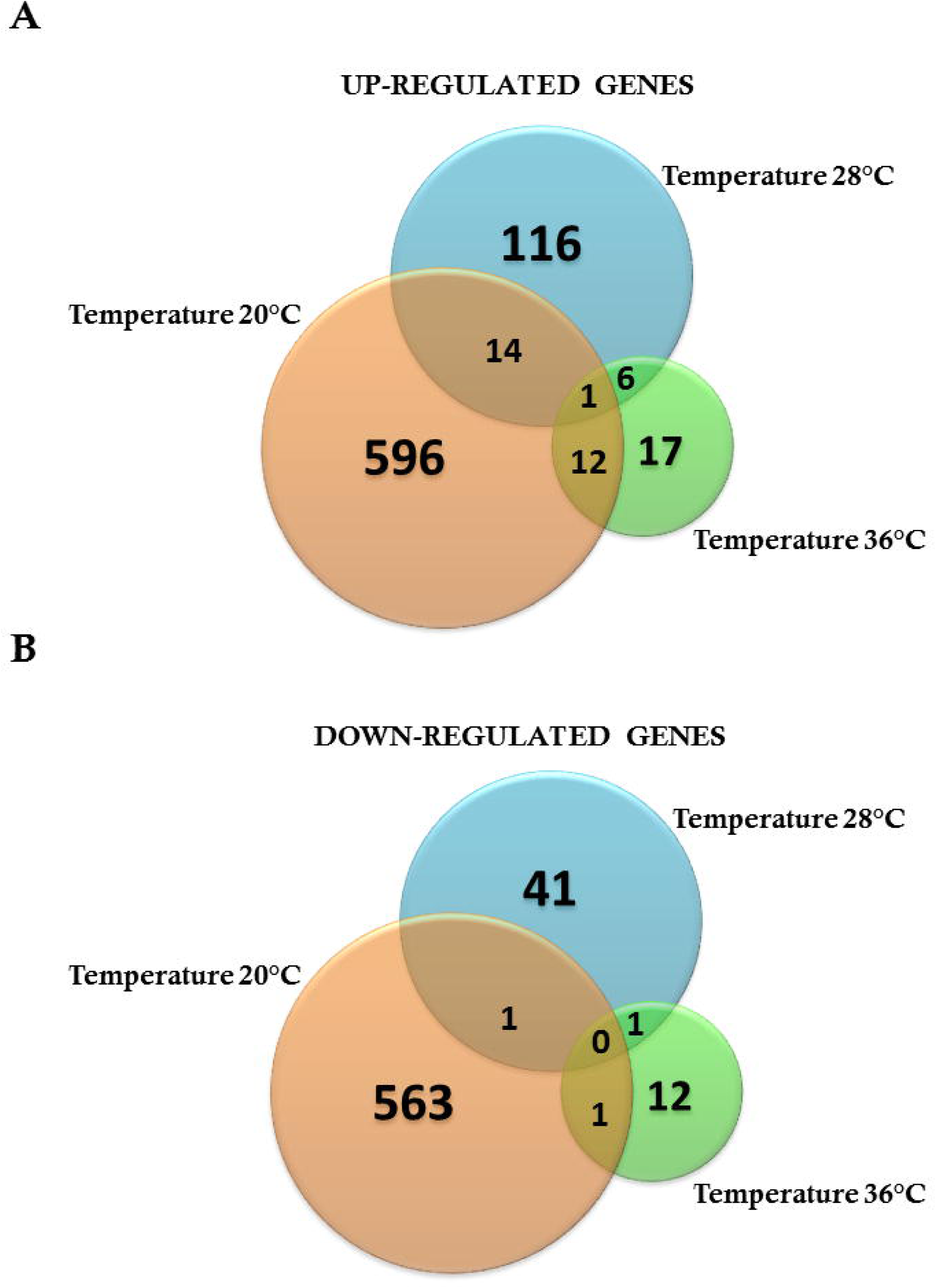
Venn diagram representing the number of specific differentially expressed genes for Zika-infected *A. aegypti* exposed in three different constant temperatures (20°C, 28°C, 36°C) (A) Up regulated genes **(B)** Down regulated genes at 48hs.

When concentrating on the top 10 most differentially expressed genes between ZIKV-exposed and unexposed mosquitoes at each temperature treatment at 48 hpf (Table 3), only at 20°C did we observe genes with altered expression (enrichment or depletion) of 10-fold or more. GO analysis demonstrated that functions associated with these genes included metallopeptidases (angiotensin converting enzyme and vitellogenic carboxypeptidase), ammonia/nitrogen metabolism (argininosuccinate lyase and alanine aminotransferase 1), and iron ion binding (a member of the cytochrome P450 family, 4g15).

**Table 3.**
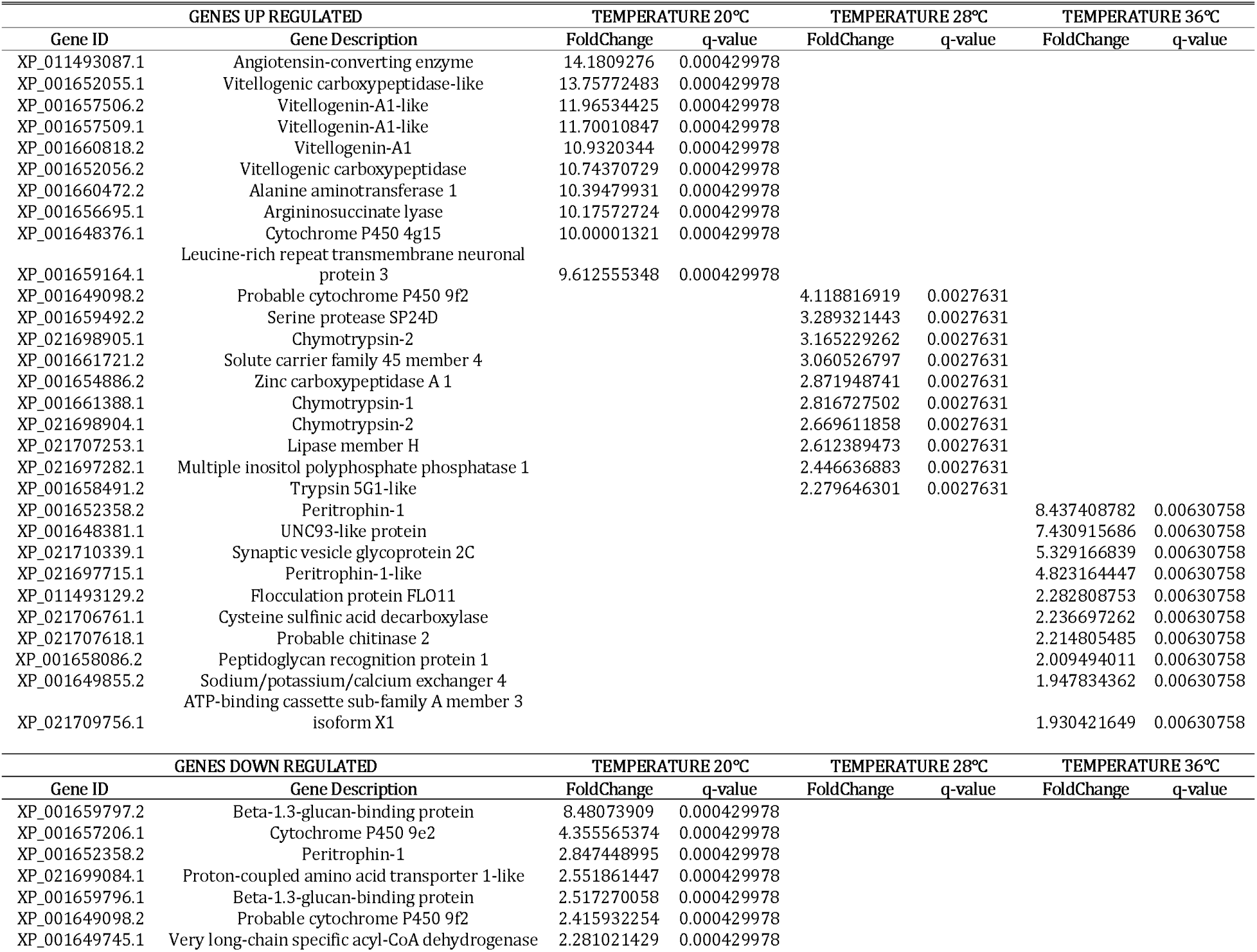
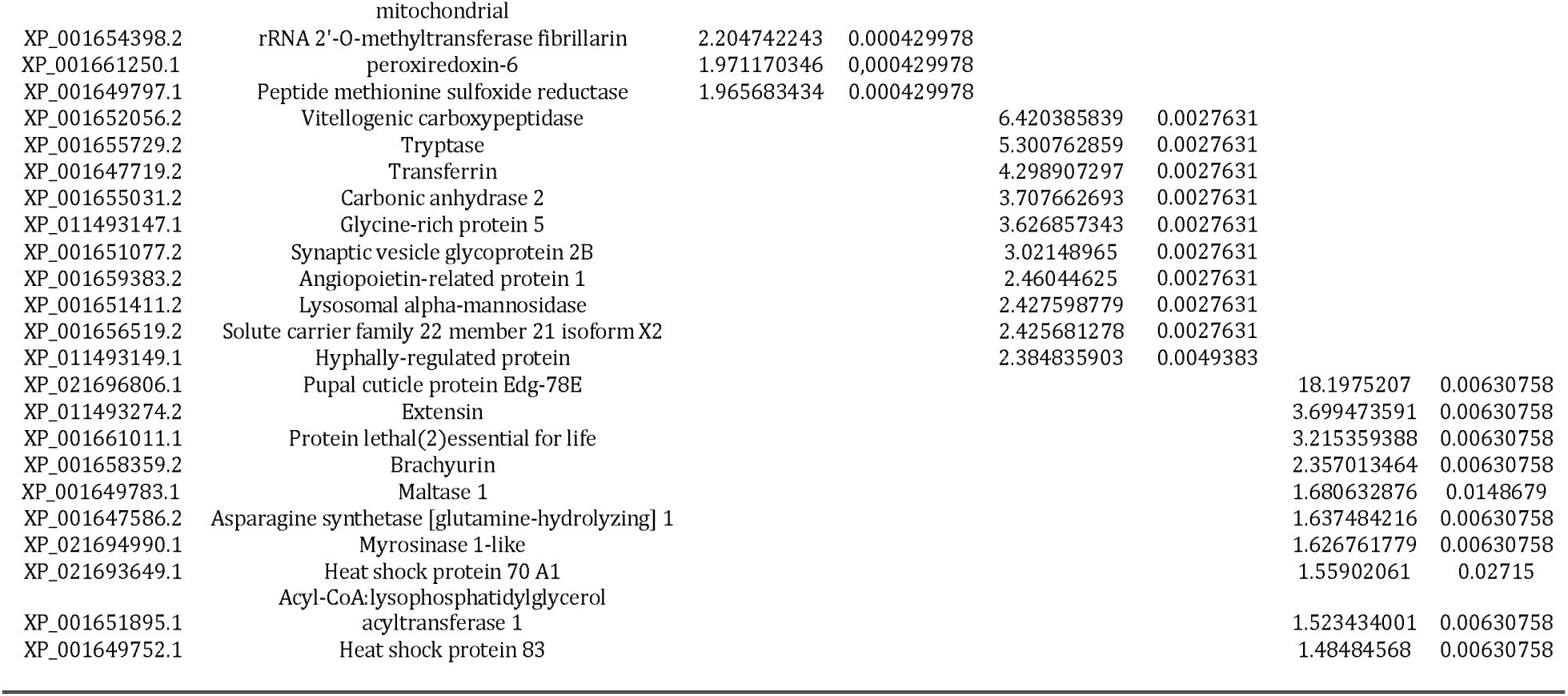
Top 10 up and down-regulated genes of ZIKV-exposed *Ae. aegypti* relative to uninfected mosquitoes within low (20°C), standard (28 °C), and high (36°C) temperatures 48 hs post-feeding blood.

Interestingly, two vitellogenins (XP_001660818.2, XP_001657509.1) that were strongly up-regulated in ZIKV-exposed and unexposed mosquitoes housed at 20°C relative to those housed at 28°C, were the genes most enriched by ZIKV exposure at the cold temperature (Table 3). The expression of these genes did not change in response to ZIKV exposure at 28°C (Data Sheet 8) and 36°C (Data Sheet 9), suggesting cold stress alters the midgut vitellogenin expression and may be more significant during a viral infection. ZIKV exposure induced a depletion of beta-1,3-glucan-binding protein (GNBP) that binds to β-1,3-glucan and lipopolysaccharide on the surface of pathogens (Dimopoulos et al., 1997) when mosquitoes were maintained at 20°C. Finally, among the most down-regulated genes in ZIKV-exposed mosquitoes at 28°C, solute carrier family 22 (XP_001656519.2), synaptic vesicle glycoprotein (XP_001651077.2) and vitellogenic carboxypeptidase (XP_001652056.2) (Table 3) were also among the most down-regulated genes in unexposed mosquitoes housed at 36°C relative to 28°C (Table 1).

## Discussion

The dynamics and distribution of vector-borne diseases depend on the interplay between the pathogen, the mosquito, and the environment. Temperature is a strong driver of vector-borne disease transmission (Kilpatrick et al., 2008; Lambrechts et al., 2011; Carrington et al., 2013a, 2013b; Mordecai et al., 2013, 2017; Johnson et al., 2015; Shocket et al., 2018; Tesla et al., 2018a). Despite the strong effects of temperature on mosquito-borne pathogens, little is known about the underpinning mechanisms (Adelman et al., 2013). In this study, RNA sequencing of *Ae. aegpyti* midguts unexposed or exposed to ZIKV early in the infection process revealed different transcriptional responses to variation in environmental temperature, with ZIKV infection modifiying these responses to temperature.

Previously we found that temperature significantly affected the efficiency of ZIKV establishing infection in *Ae. aegypti* midguts, with cool temperatures limiting ZIKV transmission primarily due to poor midgut infection, slow replication, and poor dissemination while high mosquito mortality at warmer temperatures inhibited ZIKV transmission despite efficient ZIKV infection (Tesla et al., 2018a). Similar to our previous study, we were unable to detect ZIKV replication when mosquitoes were housed in cool conditions (Figure 1), which is similar to other flavivirus systems in which infection rates were measured across different constant temperatures (Watts et al., 1987; Kilpatrick et al., 2008; Johansson et al., 2012; Xiao et al., 2014). There was no significant difference in viral RNA levels quantified from the three temperatures at 24 hpf, reflecting the initial concentration of viral RNA ingested in the blood meal (Figure 1). Therefore, we can confirm that while all treatment groups obtained ZIKV in the blood meal, only the mosquitoes housed at standard (28°C) or warm (36°C) temperatures were actively replicating virus 48 hpf.

We found that variation in temperature elicited strong expression responses in unexposed and ZIKV-exposed mosquitoes. We observed mosquitoes housed at a cold temperature (20°C) had more genes differentially expressed 48 hpf relative to mosquitoes housed under standard (28°C) and warm environments (36°C), which exhibited more similar patterns in gene expression (Figure 2A and 4A). This is not entirely surprising as metabolic theory predicts low body temperatures will inevitably depress the rates of biochemical reactions (Angilletta et al., 2010). Our data support this hypothesis, as many of the genes drastically altered at 20°C participate in blood meal digestion, peritrophic membrane (PM) formation, metabolism, and managing oxidative stress associated with the breakdown of haemoglobin into heme, a cytotoxic product (Table 1 and 2). Phosphoenolpyruvate carboxykinase and trypsin are upregulated in *Ae. aegypti* midgut during the first few hours after ingestion of a blood meal (Sanders et al., 2003), but were extraordinarily up-regulated in mosquitoes 48 hpf when housed in a cool environment in our study. Additionally, protein G12, which has previously been associated with blood meal digestion and nitrile-specific detoxification (Morlais et al., 2003; Fischer et al., 2008; Bonizzoni et al., 2011), was one of the most enriched transcripts. Further, two digestive proteases involved in glycoside hydrolysis, beta-galactosidase and alpha-N-acetylgalactosaminidase (Santamaría et al., 2015) were highly induced at 48 hpf. Glutamine synthetase, an enzyme that contributes to PM formation, was also highly induced further demonstrating that cool temperatures delay blood meal digestion (Data Sheet 1 and 3). Although the PM is semi-permeable, it is thought to form a barrier and protect the midgut from pathogens (e.g. viruses (Wang and Granados, 2000), bacteria (Kuraishi et al., 2011; Jin et al., 2019), malaria (Rodgers et al., 2017) and protozoa (Weiss et al., 2014)) and other harmful substances present in the insect gut after a blood meal (Wang et al., 2004; Shibata et al., 2015).

We also observed several genes involved in the innate immune response to be modestly upregulated in response to 20°C relative to warmer temperatures, in both unexposed and ZIKV-exposed mosquitoes. Melanization is a major effector mechanism of the mosquito immune response and has been implicated in the defense against a diversity of pathogens (e.g. bacteria (Hillyer et al., 2003a, 2003b), malaria (Kumar et al., 2003; Jaramillo-Gutierrez et al., 2009), filarial worms (Christensen et al., 2005; Huang et al., 2005), and viroses (Rodriguez-Andres et al., 2012)). Phenoloxidase, a key enzyme in the melanization pathway, was up-regulated in mosquito midguts at 20°C (Supplementary Figure S3 B and S4 B). Studies in both butterflies and *Anopheles stephensi* demonstrated that phenoloxidase activity was higher at cool temperatures and becomes less efficient at warmer temperatures (Suwanchaichinda and Paskewitz, 1998; Murdock et al., 2012b). The production of melanin is also essential for other physiological processes such as cold acclimization in insects (Crill et al., 1996; Kutch et al., 2014) the formation of the hard protective layer around eggs, and wound healing (Lai et al., 2009). Our data also reveal that c-type lectin, reported to participate in activation of the melanization cascade (Yu and Kanost, 2000; Christensen et al., 2005) was also up-regulated. Therefore, our results suggest that cold stress triggers numerous molecular changes in the mosquito, including modest changes in the levels of important immune effectors that could have important consequences for arboviral infection.

Contrary to what we observed at the cool temperature, exposure to hot conditions (36°C) does not trigger pronounced up- or down-regulation of genes relative to standard conditions (28°C). The heat shock protein 70 (HSP 70) transcript was most enriched in response to the hot environment (Table 1). The upregulation of HSP 70 is associated with reduced lifespans in other insect systems (Feder and Krebs, 1998; Feder, 1999), which may explain the rapid mosquito mortality we observed at this temperature in previous work (Tesla et al., 2018a). HSP70 has also been suggested to facilitate arbovirus infection in mosquitoes (Kuadkitkan et al., 2010; Taguwa et al., 2015).

ZIKV exposure induced very modest effects when comparing ZIKV-exposed and unexposed mosquitoes within a given temperature treatment (Figure 6, Supplementary Figure S5). This may not be entirely surprising, as ZIKV-induced transcriptional changes at standard rearing conditions (28°C) have previously been shown to be subtle 48 hr post-infection (Murdock et al., 2012a) and ZIKV was only observed to be actively replicating at 28°C and 36°C. When concentrating on differentially expressed genes between ZIKV-exposed and unexposed mosquitoes at each temperature treatment (Table 3), only at 20°C do we observe changes of 10-fold or more. However, the presence of ZIKV in the blood meal did alter the response of mosquitoes to temperature variation, with the most pronounced differences occurring in mosquitoes housed at the cool temperature. In particular, the midguts of ZIKV-exposed mosquitoes experience enhanced signal transduction processes, pH modification, midgut epithelial morphogenesis, and Toll pathway activation relative to ZIKV-exposed mosquitoes at 28°C (Figure 5), and this pattern was qualitatively different to a similar comparison in unexposed mosquitoes (Figure 3). These changes could be reflective of patterns observed in other studies that demonstrate mosquitoes infected with blood-borne pathogens actively modify ROS metabolism in midgut cells to control levels of hydrogen peroxide (H_2_O_2_), which in turn is an important modulator of downstream innate immune responses (e.g. Toll pathway), antimicrobial peptide production, and pathogen infection (Molina-Cruz et al., 2008; Herrera-Ortiz et al., 2011; Oliveira et al., 2011, 2012).

Additionally, vitellogenin proteins (Vg) were highly upregulated (>3000 fold) when ZIKV-exposed mosquitoes were housed at cool temperatures relative to ZIKV-exposed mosquitoes at 28°C (Table 2) and unexposed mosquitoes at 20°C (Table 3). Vg is precursor egg-yolk protein, but may also function by shielding cells from the negative effects of inflammation and infection (Corona et al., 2007; Azevedo et al., 2011). Work with honey bees suggest Vg-incubated insect cells display an increase in tolerance against H_2_O_2_-induced oxidative stress (Fluri et al., 1977; Seehuus et al., 2006). Vg also binds to dying cells, suggesting it may play a role in recognizing damaged cells and shielding healthy cells from toxic bi-products (Havukainen et al., 2013). Both caspase dronc (an inhibitor caspase (Cooper et al., 2007)) and an effector caspase (Bryant et al., 2008), which are important components of the apoptotic pathway controlling mechanisms of cell death, were enriched in mosquitoes housed at 20°C (Supplementary Figure S3 E and S4 E). Thus, elevated Vg expression may combat cellular damage resulting from elevated heme toxicity and oxidative stress (Havukainen et al., 2013) associated with the delayed breakdown of haemoglobin in the blood meal (Bottino-Rojas et al., 2015) in mosquitoes housed at cool temperatures. There also may be direct effect of Vg on ZIKV, as it has been associated with antiviral effects in some fish species (Garcia et al., 2010). Entry of ZIKV into mammalian cells is associated with apoptotic mimicry, with viral lipids interacting with phosphatidylserine receptors on cells gaining entry in a pathway similar to clearance of apoptotic cell (Hamel et al., 2015). Honey bee Vg binds to dying cells through interactions with lipids (Havukainen et al., 2013). Although the receptors ZIKV uses to interact and enter mosquito cells have not been identified, if Vg protein coats viral particles, it may impede normal cellular interaction and entry. Alternatively, ZIKV infection could be limited at the cooler temperature if infected mosquitoes balance ROS metabolism towards a higher state of oxidative stress as shown in other systems (Molina-Cruz et al., 2008; Herrera-Ortiz et al., 2011; Pan et al., 2012; Bottino-Rojas et al., 2015; Wong et al., 2015), facilitating downstream innate immune mechanisms and virus killing. Whether overexpression of Vg lipoproteins have direct effects on ZIKV infection or reflect a response to buffer ZIKV-exposed mosquitoes to a higher state of oxidative stress in the midgut remain open questions that will be explored in future experiments.

Finally, in ZIKV-exposed mosquitoes at 36°C, we observed peritrophin, one of the components of the peritrophic matrix, to be highly up-regulated relative to ZIKV-exposed mosquitoes housed at 28°C (Table 2) and unexposed mosquitoes housed at 36°C (Table 3). Although the PM formation is highly induced 3-24 hpf, peritrophin can undergo positive modulation by pathogens in other vector-borne disease systems (e.g. *Le. major*) (Coutinho-Abreu et al., 2013). While we cannot confirm that HSP70 or modulation of peritrophin play role in ZIKV infection, these could be potential mechanisms explaining why we detect higher viral RNA levels at this temperature in this study (Figure 1) and efficient viral dissemination and salivary gland invasion in previous work (Tesla et al., 2018a).

In this study we demonstrate profound effects of temperature on ZIKV viral replication and the transcriptional responses of mosquitoes. Temperature variation could alter the ZIKV infection process either through modifying the response of mosquitoes to ZIKV infection, altering the efficiencies of viral specific processes, or more likely both. Our study focused on midgut responses early in the infection process. However, disentangling these effects will require sampling other immunological tissues and later time points where high levels of ZIKV RNA can be detected. While further work is needed to determine the precise mechanisms at play, our results indicate that temperature shifts the balance and dynamics of the midgut environment, which could result in direct and indirect consequences for the ZIKV infection process. These results reinforce that the conventional approach of studying the mechanisms underpinning mosquito-pathogen interactions under a narrow set of laboratory conditions or across canonical innate immune pathways is likely missing important biological complexity. To move forward, we need to begin framing our mechanistic understanding of this dynamic phenotype in the ecologically variable world in which mosquitoes and pathogens associate. This study, represents a key advance towards this objective.

## Supporting information

Data Sheet 1

Data Sheet 2

Data Sheet 3

Data Sheet 4

Data Sheet 5

Data Sheet 6

Data Sheet 7

Data Sheet 8

Data Sheet 9

## Accession Numbers

The fastq raw data were deposited in NCBI SRA database under accession number SUB6324547.

## Author Contributions Statement

BT performed the experiments. PF and TM analyzed the RNAseq data. CM, MB, TM and LN designed the experiments, acquired the funding and supervised the project. All authors wrote the manuscript.

## Acknowledgements

We thank the University of Texas Medical Branch Arbovirus Reference Collection for providing the virus. We thank Dr. Americo Rodriguez from the Instituto Nacional de Salud Publica for providing mosquito eggs. We also thank the Universidade Federal de Viçosa and University of Georgia for the facilities to carry out this work. The bioinformatics analyses were performed in the UFV Cluster.

## References

Adelman, Z. N., Anderson, M. A. E., Wiley, M. R., Murreddu, M. G., Samuel, G. H., Morazzani, E. M., et al. (2013). Cooler Temperatures Destabilize RNA Interference and Increase Susceptibility of Disease Vector Mosquitoes to Viral Infection. PLoS Negl. Trop. Dis. 7, e2239. doi:10.1371/journal.pntd.0002239.

Alto, B. W., Lounibos, L. P., Mores, C. N., and Reiskind, M. H. (2008). Larval competition alters susceptibility of adult Aedes mosquitoes to dengue infection. Proc. R. Soc. B Biol. Sci. 275, 463–471. doi:10.1098/rspb.2007.1497.

Andrews, S. (2010). FastQC: a quality control tool for high throughput sequence data. Available at: http://www.bioinformatics.babraham.ac.uk/projects/fastqc.

Angilletta, M. J., Huey, R. B., and Frazier, M. R. (2010). Thermodynamic effects on organismal performance: Is hotter better? Physiol. Biochem. Zool. 83, 197–206. doi:10.1086/648567.

Azevedo, D. O., Zanuncio, J. C., Delabie, J. H. C., and Serrão, J. E. (2011). Temporal variation of vitellogenin synthesis in Ectatomma tuberculatum (Formicidae: Ectatomminae) workers. J. Insect Physiol. 57, 972–977. doi:10.1016/j.jinsphys.2011.04.015.

Barletta, A. B. F., Nascimento-Silva, M. C. L., Talyuli, O. A. C., Oliveira, J. H. M., Pereira, L. O. R., Oliveira, P. L., et al. (2017). Microbiota activates IMD pathway and limits Sindbis infection in Aedes aegypti. Parasites and Vectors. 10. doi:10.1186/s13071-017-2040-9.

Bartholomay, L. C., and Michel, K. (2018). Mosquito Immunobiology: The Intersection of Vector Health and Vector Competence. Annu. Rev. Entomol. 63, 145–167. doi:10.1146/annurev-ento-010715-023530.

Bolger, A. M., Lohse, M., and Usadel, B. (2014). Trimmomatic: A flexible trimmer for Illumina sequence data. Bioinformatics 30, 2114–2120. doi:10.1093/bioinformatics/btu170.

Bonizzoni, M., Dunn, W. A., Campbell, C. L., Olson, K. E., Dimon, M. T., Marinotti, O., et al. (2011). RNA-seq analyses of blood-induced changes in gene expression in the mosquito vector species, Aedes aegypti. BMC Genomics 12. doi:10.1186/1471-2164-12-82.

Bonizzoni, M., Dunn, W. A., Campbell, C. L., Olson, K. E., Marinotti, O., and James, A. A.. (2012). Complex Modulation of the Aedes aegypti Transcriptome in Response to Dengue Virus Infection. PLoS One 7, e50512. doi:10.1371/journal.pone.0050512.

Bottino-Rojas, V., Talyuli, O. A. C., Jupatanakul, N., Sim, S., Dimopoulos, G., Venancio, T. M., et al. (2015). Heme signaling impacts global gene expression, immunity and dengue virus infectivity in Aedes aegypti. PLoS One 10, e0135985. doi:10.1371/journal.pone.0135985.

Bryant, B., Blair, C. D., Olson, K. E., and Clem, R. J. (2008). Annotation and expression profiling of apoptosis-related genes in the yellow fever mosquito, Aedes aegypti. Insect Biochem. Mol. Biol. 38, 331–345. doi:10.1016/j.ibmb.2007.11.012.

Campbell, C. L., Keene, K. M., Brackney, D. E., Olson, K. E., Blair, C. D., Wilusz, J., et al. (2008). Aedes aegypti uses RNA interference in defense against Sindbis virus infection. BMC Microbiol. 8. doi:10.1186/1471-2180-8-47.

Carissimo, G., Pondeville, E., McFarlane, M., Dietrich, I., Mitri, C., Bischoff, E., et al. (2014). Antiviral immunity of Anopheles gambiae is highly compartmentalized, with distinct roles for RNA interference and gut microbiota. Proc. Natl. Acad. Sci. U. S. A. 112, E176–E185. doi:10.1073/pnas.1412984112.

Carrington, L. B., Armijos, M. V., Lambrechts, L., and Scott, T. W. (2013a). Fluctuations at a Low Mean Temperature Accelerate Dengue Virus Transmission by Aedes aegypti. PLoS Negl. Trop. Dis. 7, e2190. doi:10.1371/journal.pntd.0002190.

Carrington, L. B., Seifert, S. N., Armijos, M. V., Lambrechts, L., and Scott, T. W. (2013b). Reduction of Aedes aegypti vector competence for dengue virus under large temperature fluctuations. Am. J. Trop. Med. Hyg. 88, 689–697. doi:10.4269/ajtmh.12-0488.

Chauhan, C., Behura, S. K., DeBruyn, B., Lovin, D. D., Harker, B. W., Gomez-Machorro, C., et al. (2012). Comparative Expression Profiles of Midgut Genes in Dengue Virus Refractory and Susceptible Aedes aegypti across Critical Period for Virus Infection. PLoS One 7, e47350. doi:10.1371/journal.pone.0047350.

Chouin-Carneiro, T., Vega-Rua, A., Vazeille, M., Yebakima, A., Girod, R., Goindin, D., et al. (2016). Differential Susceptibilities of Aedes aegypti and Aedes albopictus from the Americas to Zika Virus. PLoS Negl. Trop. Dis. 10, 1–11. doi:10.1371/journal.pntd.0004543.

Christensen, B. M., Li, J., Chen, C. C., and Nappi, A. J. (2005). Melanization immune responses in mosquito vectors. Trends Parasitol. 21, 192–199. doi:10.1016/j.pt.2005.02.007.

Christofferson, R. C., and Mores, C. N. (2016). Potential for Extrinsic Incubation Temperature to Alter Interplay between Transmission Potential and Mortality of Dengue-Infected Aedes aegypti. Environ. Health Insights 10, 119–123. doi:10.4137/ehi.s38345.

Cirimotich, C. M., Scott, J. C., Phillips, A. T., Geiss, B. J., and Olson, K. E. (2009). Suppression of RNA interference increases alphavirus replication and virus-associated mortality in Aedes aegypti mosquitoes. BMC Microbiol. 9. doi:10.1186/1471-2180-9-49.

Cohen, J. M., Civitello, D. J., Brace, A. J., Feichtinger, E. M., Ortega, C. N., Richardson, J. C., et al. (2016). Spatial scale modulates the strength of ecological processes driving disease distributions. Proc. Natl. Acad. Sci. U. S. A. 113, E3359–E3364. doi:10.1073/pnas.1521657113.

Colpitts, T. M., Cox, J., Vanlandingham, D. L., Feitosa, F. M., Cheng, G., Kurscheid, S., et al. (2011). Alterations in the aedes aegypti transcriptome during infection with west nile, dengue and yellow fever viruses. PLoS Pathog. 7, e1002189. doi:10.1371/journal.ppat.1002189.

Conesa, A., Madrigal, P., Tarazona, S., Gomez-Cabrero, D., Cervera, A., McPherson, A., et al. (2016). A survey of best practices for RNA-seq data analysis. Genome Biol. 17. doi:10.1186/s13059-016-0881-8.

Cooper, D. M., Thi, E. P., Chamberlain, C. M., Pio, F., and Lowenberger, C. (2007). Aedes Dronc: A novel ecdysone-inducible caspase in the yellow fever mosquito, Aedes aegypti. Insect Mol. Biol. 16, 563–572. doi:10.1111/j.1365-2583.2007.00758.x.

Corona, M., Velarde, R. A., Remolina, S., Moran-Lauter, A., Wang, Y., Hughes, K. A., et al. (2007). Vitellogenin, juvenile hormone, insulin signaling, and queen honey bee longevity. Proc. Natl. Acad. Sci. U. S. A. 104, 7128–7133. doi:10.1073/pnas.0701909104.

Coutinho-Abreu, I. V., Sharma, N. K., Robles-Murguia, M., and Ramalho-Ortigao, M. (2013). Characterization of Phlebotomus papatasi Peritrophins, and the Role of PpPer1 in Leishmania major Survival in its Natural Vector. PLoS Negl. Trop. Dis. 7, e2132. doi:10.1371/journal.pntd.0002132.

Crill, W. D., Huey, R. B., and Gilchrist, G. W. (1996). Within- and Between-Generation Effects of Temperature on the Morphology and Physiology of Drosophila melanogaster. Evolution. 50, 1205–1218. doi:10.2307/2410661.

Dimopoulos, G., Richman, A., Müller, H. M., and Kafatos, F. C. (1997). Molecular immune responses of the mosquito Anopheles gambiae to bacteria and malaria parasites. Proc. Natl. Acad. Sci. U. S. A. 94, 11508–11513. doi:10.1073/pnas.94.21.11508.

Duggal, N. K., Reisen, W. K., Fang, Y., Newman, R. M., Yang, X., Ebel, G. D., et al. (2015). Genotype-specific variation in West Nile virus dispersal in California. Virology 485, 79–85. doi:10.1016/j.virol.2015.07.004.

Eng, M. W., van Zuylen, M. N., and Severson, D. W. (2016). Apoptosis-related genes control autophagy and influence DENV-2 infection in the mosquito vector, Aedes aegypti. Insect Biochem. Mol. Biol. 76, 70–83. doi:10.1016/j.ibmb.2016.07.004.

Etebari, K., Hegde, S., Saldaña, M. A., Widen, S. G., Wood, T. G., Asgari, S., et al. (2017). Global Transcriptome Analysis of Aedes aegypti Mosquitoes in Response to Zika Virus Infection. mSphere 2, e00456–17. doi:10.1128/mSphere.00456-17.

Feder, M. E. (1999). Engineering candidate genes in studies of adaptation: The heat-shock protein Hsp70 in Drosophila melanogaster. Am. Nat. 154, S55–S66.

Feder, M. E., and Krebs, R. A. (1998). Natural and genetic engineering of the heat-shock protein Hsp70 in drosophila melanogaster: Consequences for thermotolerance. Am. Zool. 38, 503–517.

Fischer, H. M., Wheat, C. W., Heckel, D. G., and Vogel, H. (2008). Evolutionary origins of a novel host plant detoxification gene in butterflies. Mol. Biol. Evol. 25, 809–820. doi:10.1093/molbev/msn014.

Fluri, P., Wille, H., Gerig, L., and Liischer, M. (1977). Juvenile hormone, vitellogenin and haemocyte composition in winter worker honeybees (Apis mellifera). Experientia 33, 1240–1241. doi:10.1007/BF01922354.

Franz, A. W. E., Kantor, A. M., Passarelli, A. L., and Clem, R. J. (2015). Tissue barriers to arbovirus infection in mosquitoes. Viruses 7, 3741–3767. doi:10.3390/v7072795.

Garcia, J., Munro, E. S., Monte, M. M., Fourrier, M. C. S., Whitelaw, J., Smail, D. A., et al. (2010). Atlantic salmon (Salmo salar L.) serum vitellogenin neutralises infectivity of infectious pancreatic necrosis virus (IPNV). Fish Shellfish Immunol. 29, 293–297. doi:10.1016/j.fsi.2010.04.010.

Girard, Y. A., Mayhew, G. F., Fuchs, J. F., Li, H., Schneider, B. S., McGee, C. E., et al. (2010). Transcriptome Changes in Culex quinquefasciatus (Diptera: Culicidae) Salivary Glands During West Nile Virus Infection. J. Med. Entomol. 47, 421–435. doi:10.1603/me09249.

Gloria-Soria, A., Armstrong, P. M., Turner, P. E., and Turner, P. E. (2017). Infection rate of aedes aegypti mosquitoes with dengue virus depends on the interaction between temperature and mosquito genotype. Proc. R. Soc. B Biol. Sci. 284. doi:10.1098/rspb.2017.1506.

Graça-Souza, A. V., Maya-Monteiro, C., Paiva-Silva, G. O., Braz, G. R. C., Paes, M. C., Sorgine, M. H. F., et al. (2006). Adaptations against heme toxicity in blood-feeding arthropods. Insect Biochem. Mol. Biol. 36, 322–335. doi:10.1016/j.ibmb.2006.01.009.

Hamel, R., Dejarnac, O., Wichit, S., Ekchariyawat, P., Neyret, A., Luplertlop, N., et al. (2015). Biology of Zika Virus Infection in Human Skin Cells. J. Virol. 89, 8880–8896. doi:10.1128/jvi.00354-15.

Havukainen, H., Münch, D., Baumann, A., Zhong, S., Halskau, Ø., Krogsgaard, M., et al. (2013). Vitellogenin recognizes cell damage through membrane binding and shields living cells from reactive oxygen species. J. Biol. Chem. 288, 28369–28381. doi:10.1074/jbc.M113.465021.

Hegde, S., Rasgon, J. L., and Hughes, G. L. (2015). The microbiome modulates arbovirus transmission in mosquitoes. Curr. Opin. Virol. 15, 97–102. doi:10.1016/j.coviro.2015.08.011.

Herrera-Ortiz, A., Martínez-Barnetche, J., Smit, N., Rodriguez, M. H., and Lanz-Mendoza, H. (2011). The effect of nitric oxide and hydrogen peroxide in the activation of the systemic immune response of Anopheles albimanus infected with Plasmodium berghei. Dev. Comp. Immunol. 35, 44–50. doi:10.1016/j.dci.2010.08.004.

Hillyer, J. F., Schmidt, S. L., and Christensen, B. M. (2003a). Hemocyte-mediated phagocytosis and melanization in the mosquito Armigeres subalbatus following immune challenge by bacteria. Cell Tissue Res. 313, 117–127. doi:10.1007/s00441-003-0744-y.

Hillyer, J. F., Schmidt, S. L., and Christensen, B. M. (2003b). Rapid Phagocytosis and Melanization of Bacteria and Plasmodium Sporozoites by Hemocytes of the Mosquito Aedes aegypti. J. Parasitol. 89, 62–69.

Huang, C. Y., Christensen, B. M., and Chen, C. C. (2005). Role of dopachrome conversion enzyme in the melanization of filarial worms in mosquitoes. Insect Mol. Biol. 14, 675–682. doi:10.1111/j.1365-2583.2005.00597.x.

Jaramillo-Gutierrez, G., Rodrigues, J., Ndikuyeze, G., Povelones, M., Molina-Cruz, A., and Barillas-Mury, C. (2009). Mosquito immune responses and compatibility between Plasmodium parasites and anopheline mosquitoes. BMC Microbiol. 9. doi:10.1186/1471-2180-9-154.

Jin, M., Liao, C., Chakrabarty, S., Wu, K., and Xiao, Y. (2019). Comparative proteomics of peritrophic matrix provides an insight into its role in Cry1Ac resistance of cotton bollworm helicoverpa armigera. Toxins. 11. doi:10.3390/toxins11020092.

Johansson, M. A., Arana-Vizcarrondo, N., Biggerstaff, B. J., Gallagher, N., Marano, N., and Staples, J. E. (2012). Assessing the risk of international spread of yellow fever virus: A mathematical analysis of an urban outbreak in Asunción, 2008. Am. J. Trop. Med. Hyg. 86, 349–358. doi:10.4269/ajtmh.2012.11-0432.

Johnson, L. R., Ben-Horin, T., Lafferty, K. D., McNally, A., Mordecai, E., Paaijmans, K. P., et al. (2015). Understanding uncertainty in temperature effects on vector-borne disease: A Bayesian approach. Ecology 96, 203–213. doi:10.1890/13-1964.1.

Kilpatrick, A. M., Meola, M. A., Moudy, R. M., and Kramer, L. D. (2008). Temperature, viral genetics, and the transmission of West Nile virus by Culex pipiens mosquitoes. PLoS Pathog. 4, e1000092. doi:10.1371/journal.ppat.1000092.

Kuadkitkan, A., Wikan, N., Fongsaran, C., and Smith, D. R. (2010). Identification and characterization of prohibitin as a receptor protein mediating DENV-2 entry into insect cells. Virology 406, 149–161. doi:10.1016/j.virol.2010.07.015.

Kumar, A., Srivastava, P., Sirisena, P. D. N. N., Dubey, S. K., Kumar, R., Shrinet, J., et al. (2018). Mosquito innate immunity. Insects 9. doi:10.3390/insects9030095.

Kumar, S., Christophides, G. K., Cantera, R., Charles, B., Han, Y. S., Meister, S., et al. (2003). The role of reactive oxygen species on Plasmodium melanotic encapsulation in Anopheles gambiae. Proc. Natl. Acad. Sci. U. S. A. 100, 14139–14144. doi:10.1073/pnas.2036262100.

Kuraishi, T., Binggeli, O., Opota, O., Buchon, N., and Lemaitre, B. (2011). Genetic evidence for a protective role of the peritrophic matrix against intestinal bacterial infection in Drosophila melanogaster. Proc. Natl. Acad. Sci. U. S. A. 108, 15966–15971. doi:10.1073/pnas.1105994108.

Kutch, I. C., Sevgili, H., Wittman, T., and Fedorka, K. M. (2014). Thermoregulatory strategy may shape immune investment in Drosophila melanogaster. J. Exp. Biol. 217, 3664–3669. doi:10.1242/jeb.106294.

Lai, S.-C., Chen, C.-C., and Hou, R. F. (2009). Immunolocalization of Prophenoloxidase in the Process of Wound Healing in the Mosquito Armigeres subalbatus (Diptera: Culicidae). J. Med. Entomol. 39, 266–274. doi:10.1603/0022-2585-39.2.266.

Lambrechts, L., Paaijmans, K. P., Fansiri, T., Carrington, L. B., Kramer, L. D., Thomas, M. B., et al. (2011). Impact of daily temperature fluctuations on dengue virus transmission by Aedes aegypti. Proc. Natl. Acad. Sci. U. S. A. 108, 7460–7465. doi:10.1073/pnas.1101377108.

Lambrechts, L., Quillery, E., Noël, V., Richardson, J. H., Jarman, R. G., Scott, T. W., et al. (2013). Specificity of resistance to dengue virus isolates is associated with genotypes of the mosquito antiviral gene Dicer-2. Proc. R. Soc. B Biol. Sci. 280. doi:10.1098/rspb.2012.2437.

Lanciotti, R. S., Kosoy, O. L., Laven, J. J., Velez, J. O., Lambert, A. J., Johnson, A. J., et al. (2008). Genetic and serologic properties of Zika virus associated with an epidemic, Yap State, Micronesia, 2007. Emerg. Infect. Dis. 14, 1232–1239. doi:10.3201/eid1408.080287.

Luckhart, S., Vodovotz, Y., Cui, L., and Rosenberg, R. (1998). The mosquito Anopheles stephensi limits malaria parasite development with inducible synthesis of nitric oxide. Proc. Natl. Acad. Sci. U. S. A. 95, 5700–5705.

Maere, S., Heymans, K., and Kuiper, M. (2005). BiNGO: A Cytoscape plugin to assess overrepresentation of Gene Ontology categories in Biological Networks. Bioinformatics 21, 3448–3449. doi:10.1093/bioinformatics/bti551.

Mccarthy, F. M., Wang, N., Magee, G. B., Nanduri, B., Lawrence, M. L., Camon, E. B., : a functional genomics resource for agriculture. BMC c Genomics 7. doi:10.1186/1471-2164-7-229.

Molina-Cruz, A., DeJong, R. J., Charles, B., Gupta, L., Kumar, S., Jaramillo-Gutierrez, G., et al. (2008). Reactive oxygen species modulate Anopheles gambiae immunity against bacteria and Plasmodium. J. Biol. Chem. 283, 3217–3223. doi:10.1074/jbc.M705873200.

Mordecai, E. A., Caldwell, J. M., Grossman, M. K., Lippi, C. A., Johnson, L. R., Neira, M., et al. (2019). Thermal biology of mosquito-borne disease. Ecol. Lett. 22, 1690–1708. doi:10.1111/ele.13335.

Mordecai, E. A., Cohen, J. M., Evans, M. V., Gudapati, P., Johnson, L. R., Lippi, C. A., et al. (2017). Detecting the impact of temperature on transmission of Zika, dengue, and chikungunya using mechanistic models. PLoS Negl. Trop. Dis. 11, e0005568. doi:10.1371/journal.pntd.0005568.

Mordecai, E. A., Paaijmans, K. P., Johnson, L. R., Balzer, C., Ben-Horin, T., de Moor, E., et al. (2013). Optimal temperature for malaria transmission is dramatically lower than previously predicted. Ecol. Lett. 16, 22–30. doi:10.1111/ele.12015.

Morlais, I., Mori, A., Schneider, J. R., and Severson, D. W. (2003). A targeted approach to the identification of candidate genes determining susceptibility to Plasmodium gallinaceum in Aedes aegypti. Mol. Genet. Genomics 269, 753–764. doi:10.1007/s00438-003-0882-7.

Murdock, C. C., Blanford, S., Hughes, G. L., Rasgon, J. L., and Thomas, M. B. (2014). Temperature alters Plasmodium blocking by Wolbachia. Sci. Rep. 4. doi:10.1038/srep03932.

Murdock, C. C., Cox-foster, D., Read, A. F., and Thomas, M. B. (2012a). Rethinking vector immunology: the role of environmental temperature in shaping resistance. Nat Rev Microbiol 10, 869–876. doi:10.1038/nrmicro2900.

Murdock, C. C., Evans, M. V., McClanahan, T. D., Miazgowicz, K. L., and Tesla, B. (2017). Fine-scale variation in microclimate across an urban landscape shapes variation in mosquito population dynamics and the potential of Aedes albopictus to transmit arboviral disease. PLoS Negl. Trop. Dis. 11. doi:10.1371/journal.pntd.0005640.

Murdock, C. C., Paaijmans, K. P., Bells, A. S., King, J. G., Hillyer, J. F., Read, A. F., et al. (2012b). Complex effects of temperature on mosquito immune function. Proc. R. Soc. B Biol. Sci. 279, 3357–3366. doi:10.1098/rspb.2012.0638.

Neill, K. O., Huang, N., Unis, D., and Clem, R. J. (2015). Rapid selection against arbovirus-induced apoptosis during infection of a mosquito vector. Proc. Natl. Acad. Sci. U. S. A. 112, E1152–E1161. doi:10.1073/pnas.1424469112.

Okech, B. A., Gouagna, L. C., Yan, G., Githure, J. I., and Beier, J. C. (2007). Larval habitats of Anopheles gambiae s.s. (Diptera: Culicidae) influences vector competence to Plasmodium falciparum parasites. Malar. J. 6. doi:10.1186/1475-2875-6-50.

Oliveira, G. de A., Lieberman, J., and Barillas-Mury, C. (2012). Epithelial Nitration by a Peroxidase/NOX5 System Mediates Mosquito Antiplasmodial Immunity. Science. 335, 856–859. doi:10.1126/science.1209678.

Oliveira, J. H. M., Gonçalves, R. L. S., Lara, F. A., Dias, F. A., Gandara, A. C. P., Menna-Barreto, R. F. S., et al. (2011). Blood meal-derived heme decreases ROS levels in the midgut of Aedes aegypti and allows proliferation of intestinal microbiota. PLoS Pathog. 7, e1001320. doi:10.1371/journal.ppat.1001320.

Pan, X., Zhou, G., Wu, J., Bian, G., Lu, P., Raikhel, A. S., et al. (2012). Wolbachia induces reactive oxygen species (ROS)-dependent activation of the Toll pathway to control dengue virus in the mosquito Aedes aegypti. Proc. Natl. Acad. Sci. U. S. A. 109, E23–E31. doi:10.1073/pnas.1116932108.

Parham, P. E., and Michael, E. (2010). Modeling the effects of weather and climate change on malaria transmission. Environ. Health Perspect. 118, 620–626. doi:10.1289/ehp.0901256.

Parham, P. E., Waldock, J., Christophides, G. K., Hemming, D., Agusto, F., Evans, K. J., et al. (2015). Climate, environmental and socio-economic change: Weighing up the balance in vector-borne disease transmission. Philos. Trans. R. Soc. B Biol. Sci. 370. doi:10.1098/rstb.2013.0551.

Rodgers, F. H., Gendrin, M., Wyer, C. A. S., and Christophides, G. K. (2017). Microbiota-induced peritrophic matrix regulates midgut homeostasis and prevents systemic infection of malaria vector mosquitoes. PLoS Pathog. 13, e1006391. doi:10.1371/journal.ppat.1006391.

Rodriguez-Andres, J., Rani, S., Varjak, M., Chase-Topping, M. E., Beck, M. H., Ferguson, M. C., et al. (2012). Phenoloxidase Activity Acts as a Mosquito Innate Immune Response against Infection with Semliki Forest Virus. PLoS Pathog. 8, e1002977. doi:10.1371/journal.ppat.1002977.

Saldaña, M. A., Etebari, K., Hart, C. E., Widen, S. G., Wood, T. G., Thangamani, S., et al. (2017a). Zika virus alters the microRNA expression profile and elicits an RNAi response in Aedes aegypti mosquitoes. PLoS Negl. Trop. Dis. 11, e0005760. doi:10.1371/journal.pntd.0005760.

Saldaña, M. A., Hegde, S., and Hughes, G. L. (2017b). Microbial control of arthropod-borne disease. Mem. Inst. Oswaldo Cruz 112, 81–93. doi:10.1590/0074-02760160373.

Sánchez-Vargas, I., Scott, J. C., Poole-Smith, B. K., Franz, A. W. E., Barbosa-Solomieu, V., Wilusz, J., et al. (2009). Dengue virus type 2 infections of Aedes aegypti are modulated by the mosquito’s RNA interference pathway. PLoS Pathog. 5, e1000299. doi:10.1371/journal.ppat.1000299.

Sanders, H. R., Evans, A. M., Ross, L. S., and Gill, S. S. (2003). Blood meal induces global changes in midgut gene expression in the disease vector, Aedes aegypti. Insect Biochem. Mol. Biol. 33, 1105–1122. doi:10.1016/S0965-1748(03)00124-3.

Sanders, H. R., Foy, B. D., Evans, A. M., Ross, L. S., Beaty, B. J., Olson, K. E., et al. (2005). Sindbis virus induces transport processes and alters expression of innate immunity pathway genes in the midgut of the disease vector, Aedes aegypti. Insect Biochem. Mol. Biol. 35, 1293–1307. doi:10.1016/j.ibmb.2005.07.006.

Santamaría, M. E., González-Cabrera, J., Martínez, M., Grbic, V., Castañera, P., Díaz, lsabel, et al. (2015). Digestive proteases in bodies and faeces of the two-spotted spider mite, Tetranychus urticae. J. Insect Physiol. 78, 69–77. doi:10.1016/j.jinsphys.2015.05.002.

Saraiva, R. G., Kang, S., Simões, M. L., Angleró-Rodríguez, Y. I., and Dimopoulos, G. (2016). Mosquito gut antiparasitic and antiviral immunity. Dev. Comp. Immunol., 53–64. doi:10.1016/j.dci.2016.01.015.

Seehuus, S.-C., Norberg, K., Gimsa, U., Krekling, T., and Amdam, G. V. (2006). Reproductive protein protects functionally sterile honey bee workers from oxidative stress. PNAS 103, 962–967. doi:10.1073/pnas.0502681103.

Shaw, W. R., and Catteruccia, F. (2019). Vector biology meets disease control: using basic research to fight vector-borne diseases. Nat. Microbiol. 4, 20–34. doi:10.1038/s41564-018-0214-7.

Shibata, T., Maki, K., Hadano, J., Fujikawa, T., Kitazaki, K., Koshiba, T., et al. (2015). Crosslinking of a Peritrophic Matrix Protein Protects Gut Epithelia from Bacterial Exotoxins. PLoS Pathog. 11. doi:10.1371/journal.ppat.1005244.

Shocket, M. S., Ryan, S. J., and Mordecai, E. A. (2018). Temperature explains broad patterns of Ross River virus transmission. Elife 7, e37762. doi:10.7554/eLife.37762.

Shragai, T., Tesla, B., Murdock, C., and Harrington, L. C. (2017). Zika and chikungunya: mosquito-borne viruses in a changing world. Ann. N. Y. Acad. Sci. 1399, 61–77. doi:10.1111/nyas.13306.

Sim, S., and Dimopoulos, G. (2010). Dengue virus inhibits immune responses in Aedes aegypti cells. PLoS One 5, e10678. doi:10.1371/journal.pone.0010678.

Simões, M. L., Caragata, E. P., and Dimopoulos, G. (2018). Diverse Host and Restriction Factors Regulate Mosquito–Pathogen Interactions. Trends Parasitol. 34, 603–616. doi:10.1016/j.pt.2018.04.011.

Siraj, A. S., Rodriguez-Barraquer, I., Barker, C. M., Tejedor-Garavito, N., Harding, D., Lorton, C., et al. (2018). Spatiotemporal incidence of Zika and associated environmental drivers for the 2015-2016 epidemic in Colombia Amir. Sci. Data. doi:10.1038/sdata.2018.73.

Souza-Neto, J. A., Sim, S., and Dimopoulos, G. (2009). An evolutionary conserved function of the JAK-STAT pathway in anti-dengue defense. Proc. Natl. Acad. Sci. U. S. A. 106, 17841–17846. doi:10.1073/pnas.0905006106.

Suwanchaichinda, C., and Paskewitz, S. M. (1998). Effects of Larval Nutrition, Adult Body Size, and Adult Temperature on the Ability of Anopheles gambiae (Diptera□: Culicidae) to Melanize Sephadex Beads. J. Med. Entomol. 35, 157–161.

Szklarczyk, D., Morris, J. H., Cook, H., Kuhn, M., Wyder, S., Simonovic, M., et al. (2017). The STRING database in 2017□: quality-controlled protein – protein association networks, made broadly accessible. Nucleic Acids Res. 45, 362–368. doi:10.1093/nar/gkw937.

Taguwa, S., Maringer, K., Li, X., Bernal-Rubio, D., Rauch, J. N., Gestwicki, J. E., et al. (2015). Defining Hsp70 Subnetworks in Dengue Virus Replication Reveals Key Vulnerability in Flavivirus Infection. Cell 163, 1108–1123. doi:10.1016/j.cell.2015.10.046.

Tchankouo-Nguetcheu, S., Khun, H., Pincet, L., Roux, P., Bahut, M., Huerre, M., et al. (2010). Differential protein modulation in midguts of Aedes aegypti infected with chikungunya and dengue 2 viruses. PLoS One 5, e13149. doi:10.1371/journal.pone.0013149.

Team, R. D. C. (2008). R: A language and environment for statistical computing. R Found. Stat. Comput. Vienna, Austria. ISBN 3-900051-07-0, URL http://www.R-project.org 3. doi:10.1007/978-3-540-74686-7.

Tesla, B., Demakovsky, L. R., Mordecai, E. A., Ryan, S. J., Bonds, M. H., Ngonghala, C. N., et al. (2018a). Temperature drives Zika virus transmission: Evidence from empirical and mathematical models. Proc. R. Soc. B Biol. Sci. 285. doi:10.1098/rspb.2018.0795.

Tesla, B., Demakovsky, L. R., Packiam, H. S., Mordecai, E. A., Rodríguez, A. D., Bonds, M. H., et al. (2018b). Estimating the effects of variation in viremia on mosquito susceptibility, infectiousness, and R0 of Zika in Aedes aegypti. PLoS Negl. Trop. Dis. 12, e0006733. doi:10.1371/journal.pntd.0006733.

Trapnell, C., Pachter, L., and Salzberg, S. L. (2009). TopHat: Discovering splice junctions with RNA-Seq. Bioinformatics 25, 1105–1111. doi:10.1093/bioinformatics/btp120.

Trapnell, C., Roberts, A., Goff, L., Pertea, G., Kim, D., Kelley, D. R., et al. (2013). Differential gene and transcript expression analysis of RNA-seq experiments with TopHat and Cufflinks. Nat. Protoc. 7, 562–578. doi:10.1038/nprot.2012.016.

Wang, H., Gort, T., Boyle, D. L., and Clem, R. J. (2012). Effects of Manipulating Apoptosis on Sindbis Virus Infection of Aedes aegypti Mosquitoes. J. Virol. 86, 6546–6554. doi:10.1128/jvi.00125-12.

Wang, P., and Granados, R. R. (2000). Calcofluor disrupts the midgut defense system in insects. Insect Biochem. Mol. Biol. 30, 135–143. doi:10.1016/S0965-1748(99)00108-3.

Wang, P., Li, G., and Granados, R. R. (2004). Identification of two new peritrophic membrane proteins from larval Trichoplusia ni: Structural characteristics and their functions in the protease rich insect gut. Insect Biochem. Mol. Biol. 34, 215–227. doi:10.1016/j.ibmb.2003.10.001.

Watts, D. M., Burke, D. S., Harrison, B. A., Whitmire, R. E., and Nisalak, A. (1987). Effect of Temperature on the Vector Efficiency of Aedes aegypti for Dengue 2 Virus. Am. J. Trop. Med. Hyg. 36, 143–152.

Weiss, B. L., Savage, A. F., Griffith, B. C., Wu, Y., and Aksoy, S. (2014). The Peritrophic Matrix Mediates Differential Infection Outcomes in the Tsetse Fly Gut following Challenge with Commensal, Pathogenic, and Parasitic Microbes. J. Immunol. 193, 773–782. doi:10.4049/jimmunol.1400163.

Wilke, A. B. B., and Marrelli, M. T. (2015). Paratransgenesis: A promising new strategy for mosquito vector control. Parasites and Vectors 8. doi:10.1186/s13071-015-0959-2.

Willard, K. A., Demakovsky, L., Tesla, B., Goodfellow, F. T., Stice, S. L., Murdock, C. C., et al. (2017). Zika virus exhibits lineage-specific phenotypes in cell culture, in Aedes aegypti mosquitoes, and in an embryo model. Viruses 9. doi:10.3390/v9120383.

Willard, K. A., Elling, C. L., Stice, S. L., and Brindley, M. A. (2019). The oxysterol 7-ketocholesterol reduces zika virus titers in vero cells and human neurons. Viruses 11, 1–17. doi:10.3390/v11010020.

Wong, Z. S., Brownlie, J. C., and Johnson, K. N. (2015). Oxidative stress correlates with Wolbachia-mediated antiviral protection in Wolbachia-Drosophila associations. Appl. Environ. Microbiol. 81, 3001–3005. doi:10.1128/AEM.03847-14.

Xi, Z., Ramirez, J. L., and Dimopoulos, G. (2008). The Aedes aegypti toll pathway controls dengue virus infection. PLoS Pathog. 4, e1000098. doi:10.1371/journal.ppat.1000098.

Xiao, F. Z., Zhang, Y., Deng, Y. Q., He, S., Xie, H. G., Zhou, X. N., et al. (2014). The effect of temperature on the extrinsic incubation period and infection rate of dengue virus serotype 2 infection in Aedes albopictus. Arch. Virol. 159, 3053–3057. doi:10.1007/s00705-014-2051-1.

Yen, P.-S., James, A., Li, J.-C., Chen, C.-H., and Failloux, A.-B. (2018). Synthetic miRNAs induce dual arboviral-resistance phenotypes in the vector mosquito Aedes aegypti. *Commun*. Biol. 1. doi:10.1038/s42003-017-0011-5.

Yu, X. Q., and Kanost, M. R. (2000). Immulectin-2, a lipopolysaccharide-specific lectin from an insect, Manduca sexta, is induced in response to Gram-negative bacteria. J. Biol. Chem. 275, 37373–37381. doi:10.1074/jbc.M003021200.

Zouache, K., Fontaine, A., Vega-Rua, A., Mousson, L., Thiberge, J. M., Lourenco-De-Oliveira, R., et al. (2014). Three-way interactions between mosquito population, viral strain and temperature underlying chikungunya virus transmission potential. Proc. R. Soc. B Biol. Sci. 281. doi:10.1098/rspb.2014.1078.

