## Supplementary material for "Temperature Dramatically Shapes Mosquito Gene Expression with Consequences for Mosquito-Zika Virus Interactions": SI Figures and Legends

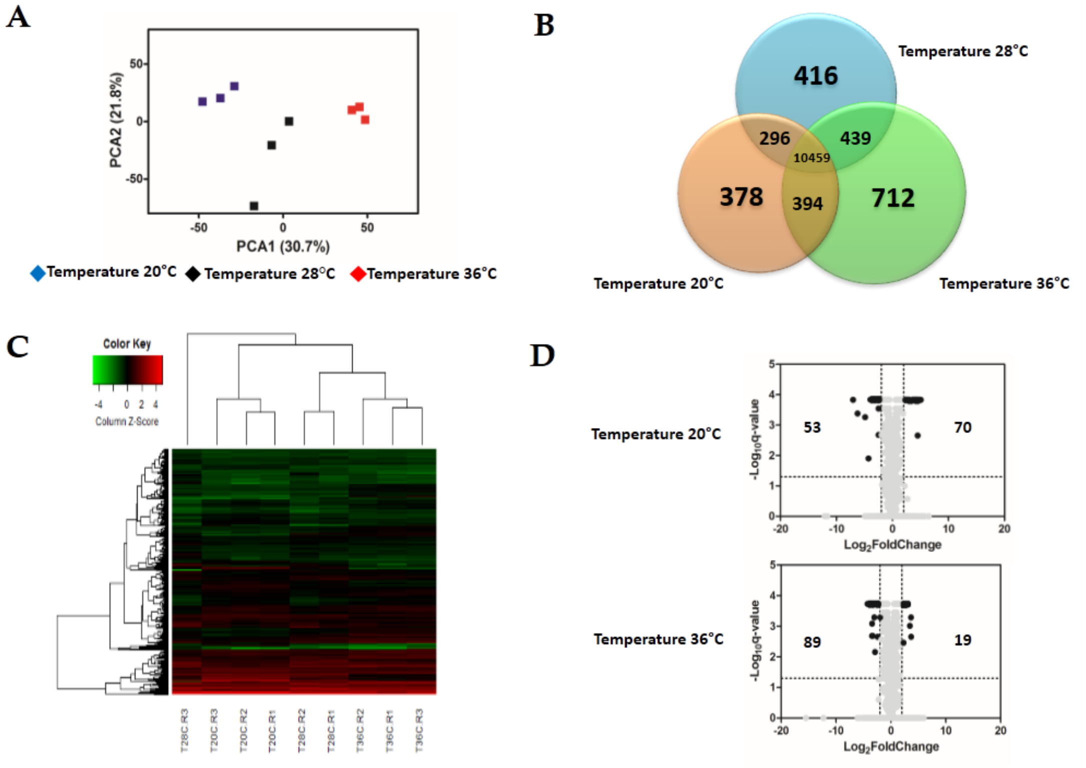
A
PCA2 (21.8%)
PCA1 (30.7%)
Temperature 20°C
Temperature 28°C
Temperature 36°C
B
Temperature 28°C
416
296
439
10459
378
394
712
Temperature 20°C
Temperature 36°C
C
Color Key
Column Z-Score
T28C R3
T20C R3
T20C R2
T20C R1
T28C R2
T28C R1
T36C R2
T36C R1
T36C R3
D
Temperature 20°C
53
70
Temperature 36°C
89
19
Log2FoldChange
-Log10q-value

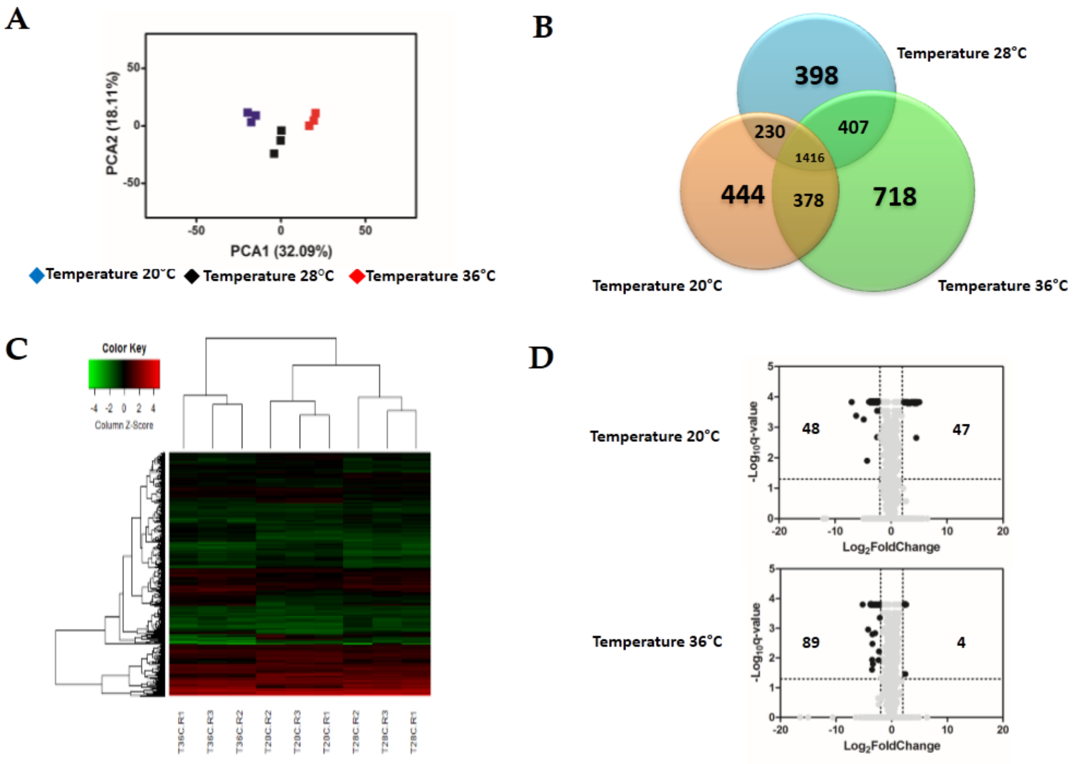

A
PCA2 (18.11%)
PCA1 (32.09%)
Temperature 20°C
Temperature 28°C
Temperature 36°C
B
Temperature 28°C
398
230
407
1416
444
378
718
Temperature 20°C
Temperature 36°C
C
Color Key
Column Z-Score
T36C R1
T36C R3
T36C R2
T20C R2
T20C R3
T20C R1
T28C R2
T28C R3
T28C R1
D
Temperature 20°C
48
47
-Log10q-value
Log2FoldChange
Temperature 36°C
89
4
-Log10q-value
Log2FoldChange

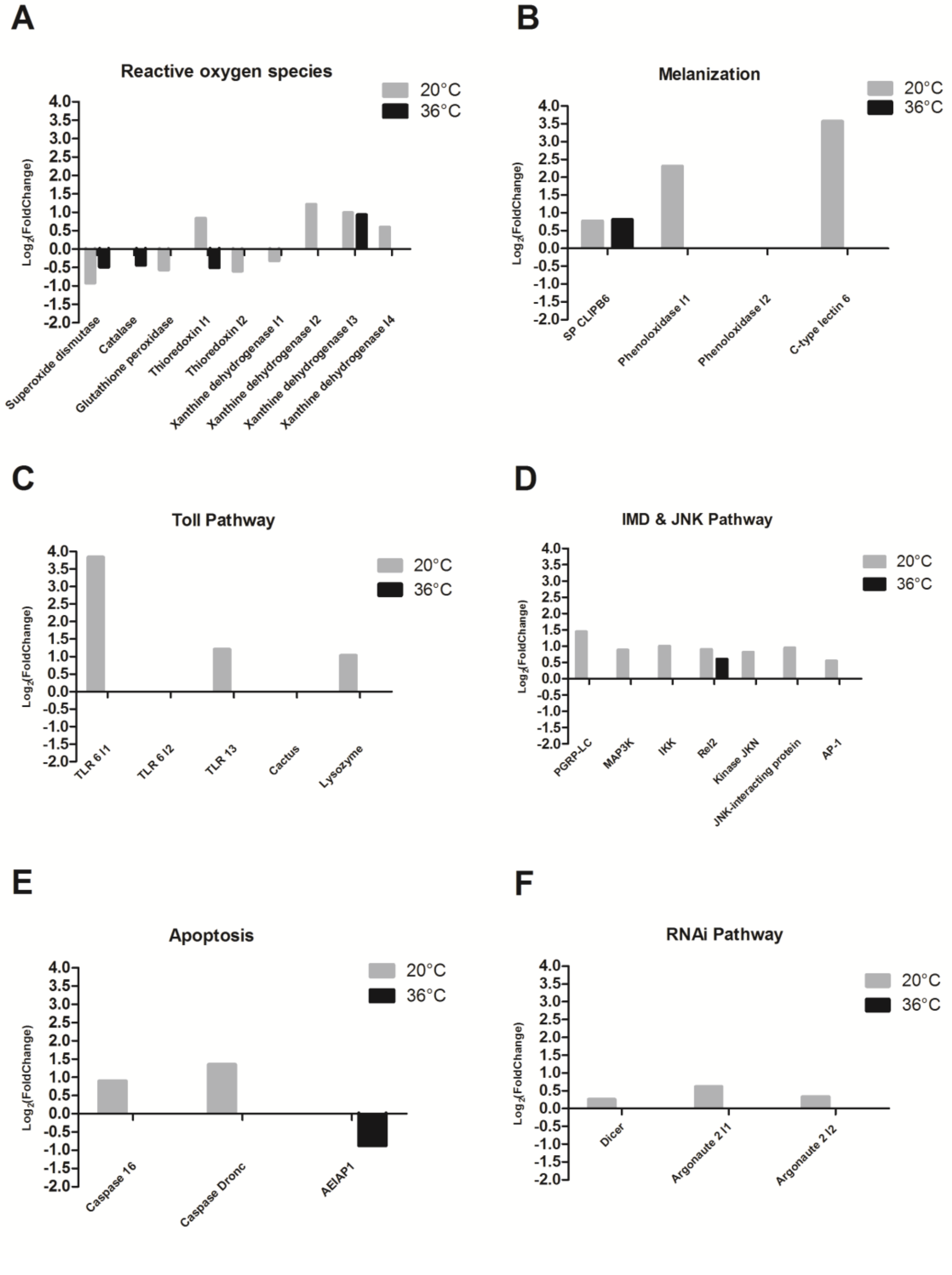
A
Reactive oxygen species
20°C
36°C
Log2(FoldChange)
4.0
3.5
3.0
2.5
2.0
1.5
1.0
0.5
0.0
-0.5
-1.0
-1.5
-2.0
Superoxide dismutase
Catalase
Glutathione peroxidase
Thioredoxin I1
Thioredoxin I2
Xanthine dehydrogenase I1
Xanthine dehydrogenase I2
Xanthine dehydrogenase I3
Xanthine dehydrogenase I4
B
Melanization
20°C
36°C
Log2(FoldChange)
SP CLIPB6
Phenoloxidase I1
Phenoloxidase I2
C-type lectin 6
C
Toll Pathway
20°C
36°C
Log2(FoldChange)
TLR 6 I1
TLR 6 I2
TLR 13
Cactus
Lysozyme
D
IMD & JNK Pathway
20°C
36°C
Log2(FoldChange)
PGRP-LC
MAP3K
IKK
Rel2
Kinase JKN
JNK-interacting protein
AP-1
E
Apoptosis
20°C
36°C
Log2(FoldChange)
Caspase 16
Caspase Dronc
AEIAP1
F
RNAi Pathway
20°C
36°C
Log2(FoldChange)
Dicer
Argonaute 2 I1
Argonaute 2 I2

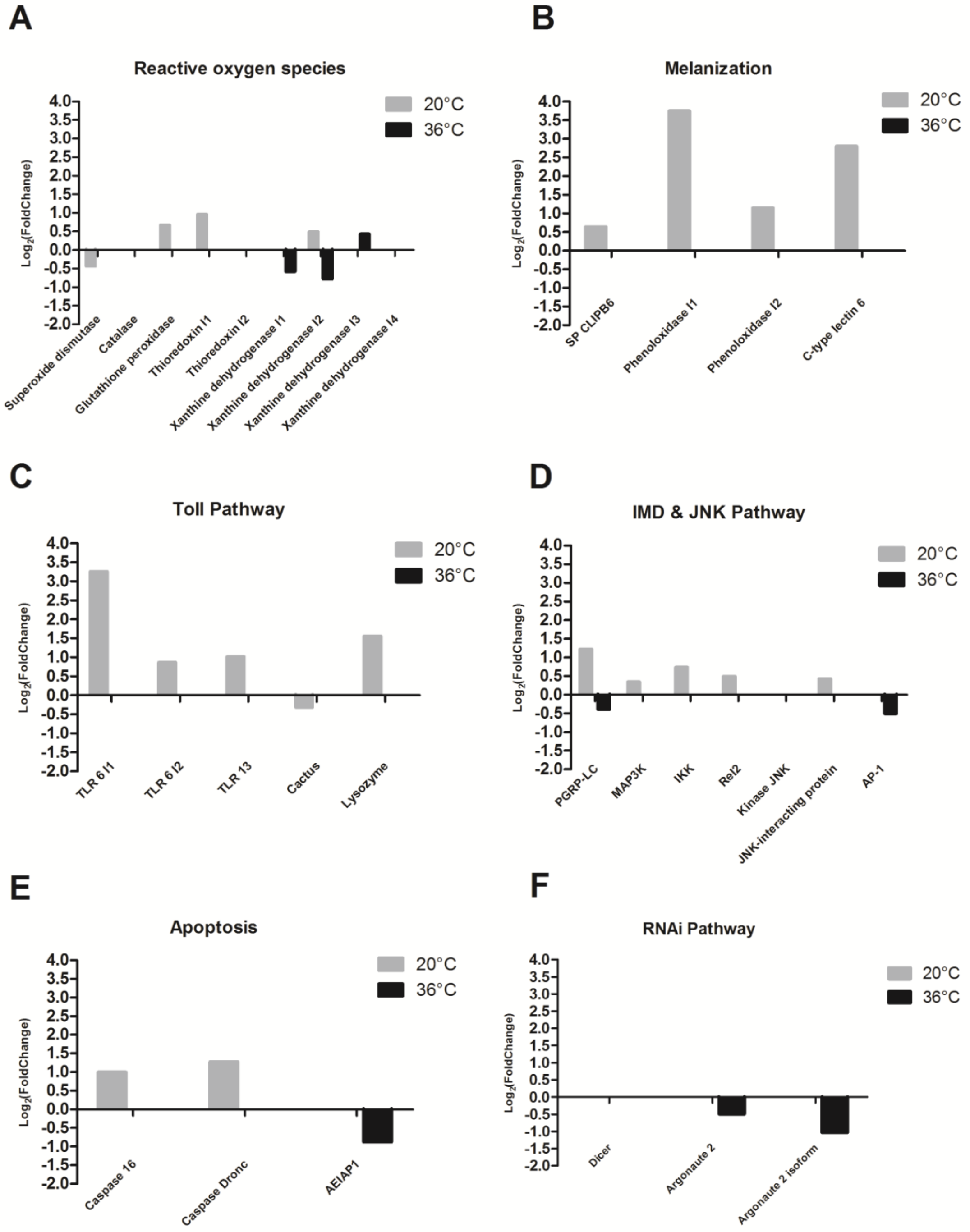
A
Reactive oxygen species
20°C
36°C
Log2(FoldChange)
Superoxide dismutase
Catalase
Glutathione peroxidase
Thioredoxin I1
Thioredoxin I2
Xanthine dehydrogenase I1
Xanthine dehydrogenase I2
Xanthine dehydrogenase I3
Xanthine dehydrogenase I4
B
Melanization
SP CLIPB6
Phenoloxidase I1
Phenoloxidase I2
C-type lectin 6
C
Toll Pathway
TLR 6 I1
TLR 6 I2
TLR 13
Cactus
Lysozyme
D
IMD & JNK Pathway
PGRP-LC
MAP3K
IKK
Rel2
Kinase JNK
JNK-interacting protein
AP-1
E
Apoptosis
Caspase 16
Caspase Dronc
AEIAP1
F
RNAi Pathway
Dicer
Argonaute 2
Argonaute 2 isoform

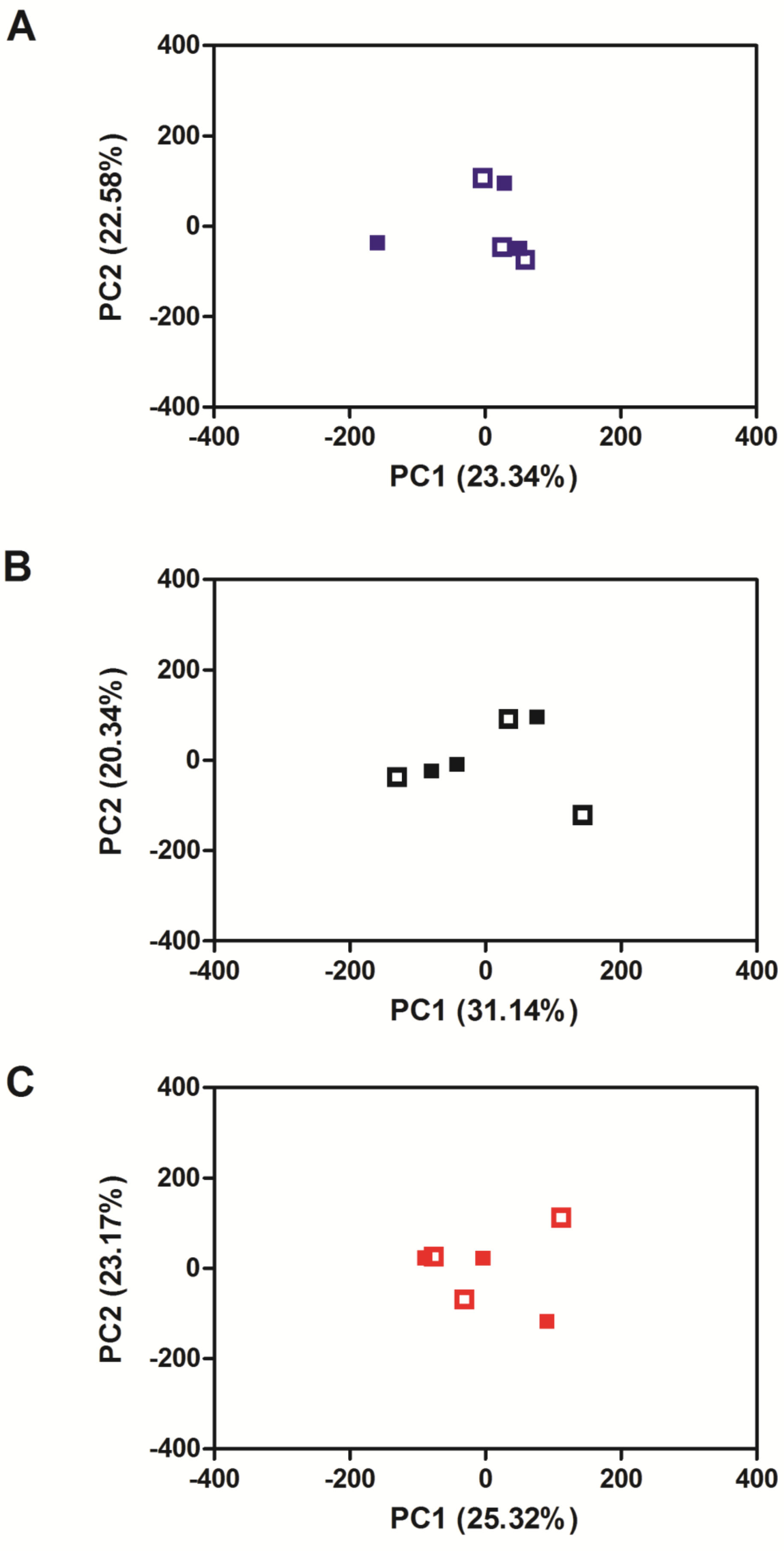
A
B
C
PC1 (23.34%)
PC2 (22.58%)
PC1 (31.14%)
PC2 (20.34%)
PC1 (25.32%)
PC2 (23.17%)
400
200
0
-200
-400

A

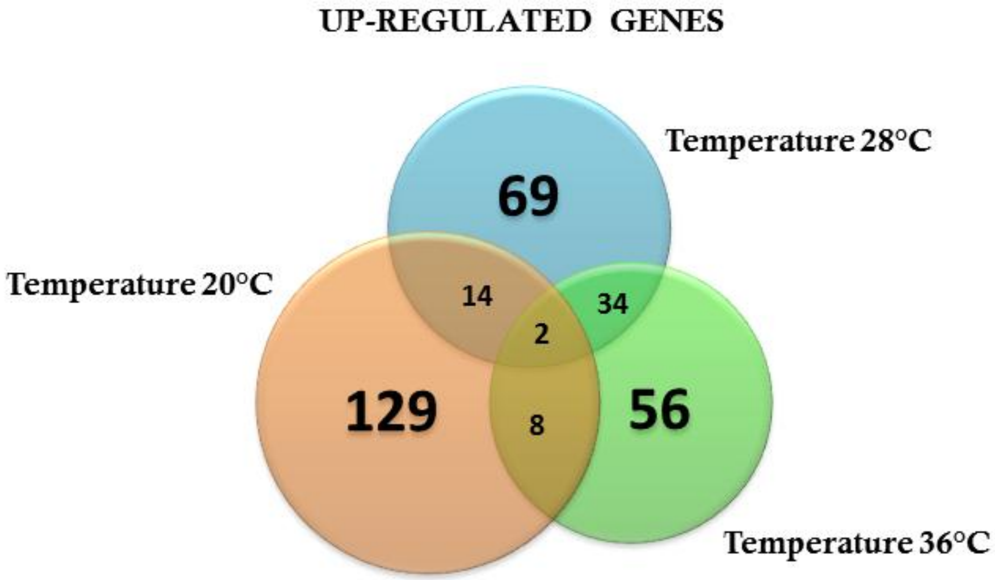
UP-REGULATED GENES
Temperature 28°C
Temperature 20°C
Temperature 36°C
69
14
34
2
129
8
56

B

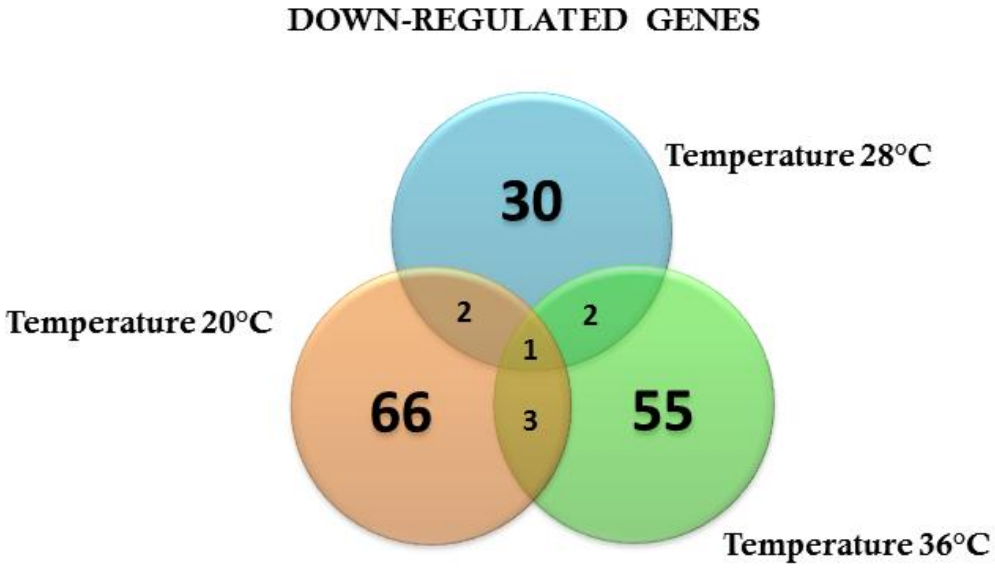
DOWN-REGULATED GENES
Temperature 28°C
Temperature 20°C
Temperature 36°C
30
2
2
1
66
3
55

**Supplementary Material Captions**

**Supplementary Figure S1. Effect of the temperature in the expression profile of non-infected *A. aegypti* (A)** Principal Component Analysis (PCA) plot showing the global gene expression profiles **(B)** Heatmap plot showing local differences (TC20R1: represents the replicate 1 for the temperature 20ºC) **(C)** Venn Diagram reporting the number of specific and shared genes **(D)** Volcano plot representing the differential gene expression from RNAseq samples from *Aedes aegypti* exposed in three different constant temperatures (20ºC, 28ºC, 36ºC) at 24 hs.

**Supplementary Figure S2. Effect of the temperature in the expression profile of Zika-infected *A. aegypti* (A)** Principal Component Analysis (PCA) plot showing the global gene expression profiles **(B)** Heatmap plot showing local differences (TC20R1: represents the replicate 1 for the temperature 20ºC) **(C)** Venn Diagram reporting the number of specific and shared genes **(D)** Volcano plot representing the differential gene expression from RNAseq samples from infected *Aedes aegypti* exposed in three different constant temperatures (20ºC, 28ºC, 36ºC) at 24 hs.

**Supplementary Figure S3. Analysis of differentially expressed genes related to the immune response in uninfected mosquitoes maintained at 20°C and 36°C compared to those kept at 28°C 48 hours after blood-feeding (A)** Reactive Oxygen Species; **(B)** Melanization; **(C)** Toll Pathway; **(D)** IMD & JNK Pathway; **(E)** Apoptosis; **(F)** RNAi Pathway.

**Supplementary Figure S4. Analysis of differentially expressed genes related to the immune response in ZIKV-infected mosquitoes maintained at 20°C and 36°C compared to those kept at 28°C 48 hours after blood-feeding (A)** Reactive Oxygen Species; **(B)** Melanization; **(C)** Toll Pathway; **(D)** IMD & JNK Pathway; **(E)** Apoptosis; **(F)** RNAi Pathway.

**Supplementary Figure S5. The Principal Component Analysis (PCA) of the effect of the Zika infection in three different constant temperatures (A) 20ºC (B) 28ºC (C) 36ºC at 24hs.**

**Supplementary Figure S6. Venn diagram representing the number of specific differentially expressed genes for Zika infected *A. aegypti* exposed in three different constant temperatures (20ºC, 28ºC, 36ºC) (A)** Up regulated genes **(B)** Down regulated genes at 24hs.

**Data Sheet 1. Differentially expressed genes in mosquitoes held at 20°C relative to those held at 28°C 24 and 48hs after uninfected control blood meal.**

**Data Sheet 2. Differentially expressed genes in mosquitoes held at 36°C relative to those held at 28°C 24 and 48hs after uninfected control blood meal.**

**Data Sheet 3. Differentially expressed genes involved in oxidative stress and innate immunity in mosquitoes held at 20°C and 36°C relative to those held at 28°C 48hs after uninfected control blood meal.**

**Data Sheet 4. Differentially expressed genes involved in oxidative stress and innate immunity in mosquitoes held at 20°C and 36°C relative to those held at 28°C 48hs after ZIKV-positive blood meal.**

**Data Sheet 5. Differentially expressed genes in mosquitoes held at 20°C relative to those held at 28°C 24 and 48hs after ZIKV-positive blood meal.**

**Data Sheet 6. Differentially expressed genes in mosquitoes held at 36°C relative to those held at 28°C 24 and 48hs after ZIKV-positive blood meal.**

**Data Sheet 7. Differentially expressed genes in mosquitoes 24 and 48 hours after ZIKV-positive blood meal relative to mosquitoes fed with uninfected blood held at 20°C.**

**Data Sheet 8. Differentially expressed genes in mosquitoes 24 and 48 hours after ZIKV-positive blood meal relative to mosquitoes fed with uninfected blood held at 28°C.**

**Data Sheet 9. Differentially expressed genes in mosquitoes 24 and 48 hours after**

**ZIKV-positive blood meal relative to mosquitoes fed with uninfected blood held at 36°C.**
